# A synthetic signalling network imitating the action of immune cells in response to bacterial metabolism

**DOI:** 10.1101/2023.02.02.526524

**Authors:** Michal Walczak, Leonardo Mancini, Jiayi Xu, Federica Raguseo, Jurij Kotar, Pietro Cicuta, Lorenzo Di Michele

## Abstract

State-of-the-art bottom-up synthetic biology allows us to replicate many basic biological functions in artificial cell-like devices. To mimic more complex behaviours, however, *artificial cells* would need to perform many of these functions in a synergistic and coordinated fashion, which remains elusive. Here we considered a sophisticated biological response, namely the capture and deactivation of pathogens by neutrophil immune cells, through the process of netosis. We designed a consortium consisting of two synthetic agents – responsive DNA-based particles and antibiotic-loaded lipid vesicles – whose coordinated action mimics the sought immune-like response when triggered by bacterial metabolism. The artificial netosis-like response emerges from a series of interlinked sensing and communication pathways between the live and synthetic agents, and translates into both physical and chemical antimicrobial actions, namely bacteria immobilisation and exposure to antibiotics. Our results demonstrate how advanced life-like responses can be prescribed with a relatively small number of synthetic molecular components, and outlines a new strategy for artificial-cell-based antimicrobial solutions.

## INTRODUCTION

Recent years have witnessed a substantial growth in the fields of artificial cell science and bottom-up synthetic biology [1–6], which aim at producing cell mimics capable of replicating the behaviours of biological cells. *Artificial cells* hold applicative potential in diagnostics [7] and therapeutics [8–11], as well as in fundamental biology and [12] origin of life research [13]. This interest has led to a rapid expansion of the available design toolkit for artificial cells, enabling replication of biological processes and features such as division [14–16], metabolism [17, 18], growth [19], motility [20, 21], communication [22, 23], sensing [24, 25], compartmentalisation [26–28], protein expression [29] and DNA replication [30]. Artificial cells have been successfully built from membrane-bound scaffolds in the form of liposomes [31, 32], polymersomes [31, 33] and proteinosomes [34, 35], as well as from membrane-less coacervates or hydrogels [27, 36–39].

While these early successes in mimicking individual biological functions carry both applicative and fundamental interest, challenges remain in place when attempting to design artificial cells capable of sustaining more advanced biomimetic responses, where many elementary functions need to be performed in a coordinated way. Among the advanced behaviours we seek to implement are those that resemble the action of the immune system [40, 41], which would unlock disruptive applications of artificial cells to *in vivo* therapeutics [42].

In this scenario, artificial cells would need to establish multi-agent signalling and signal transduction networks with biological and other artificial cells, involving several interlinked responses: *i)* Detection and transduction of signals generated by live cells; *ii)* Communication with live cells and possibly other artificial cells; *iii)* Information processing; *iv)* Individual and/or collective responses that may involve both physical and chemical action, *e*.*g*. the release of therapeutic agents and the mechanical perturbation of live cells.

Successful attempts at establishing signalling networks between live and artificial cells have been reported, often involving one-way communication, *e*.*g*. triggering of bacterial gene expression [43–50], and more rarely twoway pathways leading to cell death [10]. However, these remarkable examples are still relatively simple, relying on one or a small number of individual functionalities, and thus failing to address the need to engineer more advanced emerging behaviours.

In this contribution, we aim to build a synthetic signalling network that mimics a complex response of the innate immune system, namely that of *netosis* [51–53], whereby neutrophils excrete a sticky *Neutrophil Extracellular Trap* (NET) formed from their genomic DNA, which traps pathogens and then disrupts them thanks to embedded antimicrobial proteins (Fig. 1**a**). As sketched in Fig. 1**b**, the synthetic pathway we propose features two artificial cell-like agents, *i)* responsive DNA-based particles [54] and *ii)* antibiotic-loaded liposomes, whose coordinated action, triggered by bacterial activity, gives rise to the sought behaviour. The DNA-particles sense a decrease in pH resulting from the natural glucose metabolism of *E. coli* [55], and respond by exposing their hydrophobic core, which then leads to the formation of a *synthetic DNA NET*. The sticky material traps and immobilises the bacteria and, at the same time, permeabilises the liposomes, which ultimately release an antibiotic able to hinder growth of the trapped cells. The synthetic netosis pathway coordinates several functionalities, including one-way (particles → liposomes and liposomes → bacteria) and two-way (particles ↔ bacteria) communication, a collective self-assembly response, and both physical and chemical interference with bacterial activity. Our proof-of-concept implementation thus demonstrates how advanced life-like behaviours can be engineered from the bottom-up, relying on a relatively small number of molecular and nanoscale components. In addition, the platform represents a starting point for the development of biomimetic antimicrobial solutions and, more generally, synthetic-cell therapeutics.

**FIG. 1.**
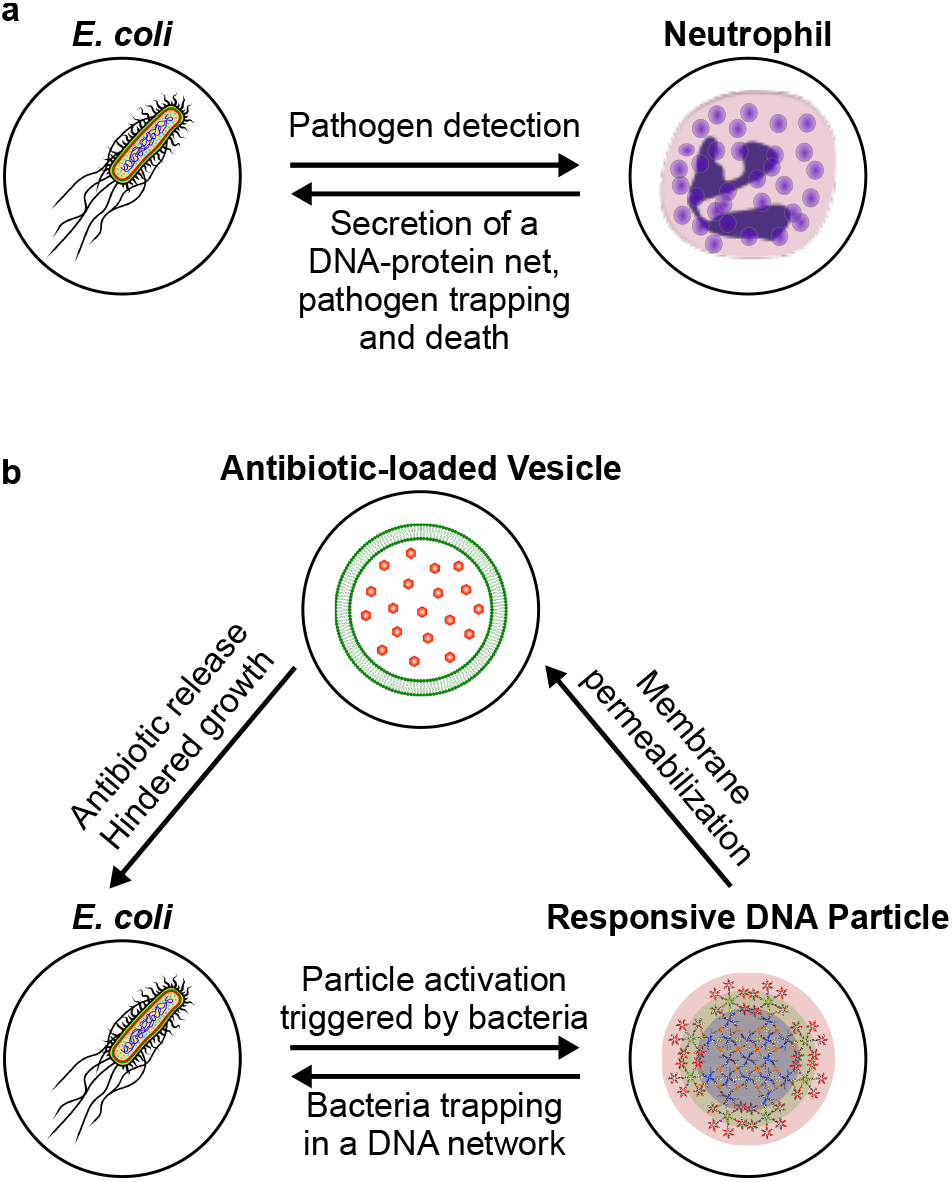
Three-agent synthetic consortium mimics the antimicrobial response of innate immune cells. **a**. Neutrophils respond to the presence of pathogens by excreting a mesh formed by their genomic DNA, histones, and granules containing molecules with antimicrobial properties. The Neutrophil Extracellular Trap (NET) immobilises the pathogens and kills them due to the antimicrobial properties of the molecules associated. **b**. Three-agent synthetic consortium designed to exhibit a netosis-like response. Responsive DNA-based particles detect model targets (*E. coli*) by sensing medium acidification induced by the glucose metabolism of the cells. “Activated” particles respond by forming a sticky DNA-cholesterol network that immobilises the bacteria and, at the same time, permeabilises antibiotic-loaded Giant Unilamellar Vesicles (GUVs). Antibiotic release suppresses bacterial growth and division, fulfilling the antimicrobial role of the DNA-binding proteins in the biological system.

## RESULTS

### Fabrication of pH-responsive DNA particles

Figure 2**a** shows the DNA-based particles used in this work, which feature a core-shell structure with an hydrophobised centre surrounded by an hydrophilic outer layer, the latter ensuring colloidal stability. The amphiphilic *core motifs* (CM) consist of 4-pointed DNA junctions (nanostars), with the end of each double-stranded (ds) DNA arm labelled by a cholesterol moiety. Similar “C-star” designs have been shown to self-assemble into framework-like materials sustained by cholesterol-mediated hydrophobic forces, with programmable nanoscale morphology and multi-stimuli-responsiveness [27, 54, 61–63]. The shell comprises of two 6-pointed all-DNA nanostars, labelled as *inner* and *outer corona motifs* (ICM and OCM, respectively), together forming dendrimeric construct that can connect to core motifs through single-stranded (ss) DNA overhangs (Fig. 2**a**). As we recently demonstrated, the core-shell particle morphology can be attained following a two-step thermal annealing process, which can be adapted to prescribe the size of the particles, from a few hundred nanometres to several microns [54]. Figure 2**b** shows a large particle where the sought core-shell structure is clearly visible in confocal microscopy. Particles of this large size were purposefully created with a modified assembly protocol to enable facile confocal visualisation. For the remainder of this work, the assembly protocol was set up to produce smaller particles, with hydrodynamic radius of either ∼ 1 *µ*m, ∼ 750 nm, or ∼ 200 nm, depending on the specific experiment. Details of sample preparation are provided in the Methods, while sequences of all oligonucleotides used in the work and composition of all samples are shown in Supplementary Tables 1 and 2, respectively. Correct assembly of the individual core and corona motifs was verified with both Agarose Gel Electrophoresis (AGE) and Dynamic Light Scattering (DLS), summarised in Supplementary Figs 1 and 2, while their thermal stability was assessed with UV-absorbance (Supplementary Fig. 3). In our previous contribution, we have demonstrated that removal of the corona, induced by a non-biologically-relevant ssDNA trigger, leads to exposure of the particles’ hydrophobic cores and formation of a sticky DNA-cholesterol network [54]. The network has then been shown to trap motile *E. coli* and, independently, disrupt Giant Unilamellar lipid Vesicles (GUVs) [54]. These are two key functionalities for the netosis-like signalling network we seek to implement here (Fig. 1**b**). However, multiple critical features are missing from the system presented in Ref. [54], including the ability of the particles to directly respond to signals generated by the target cells.

**FIG. 2.**
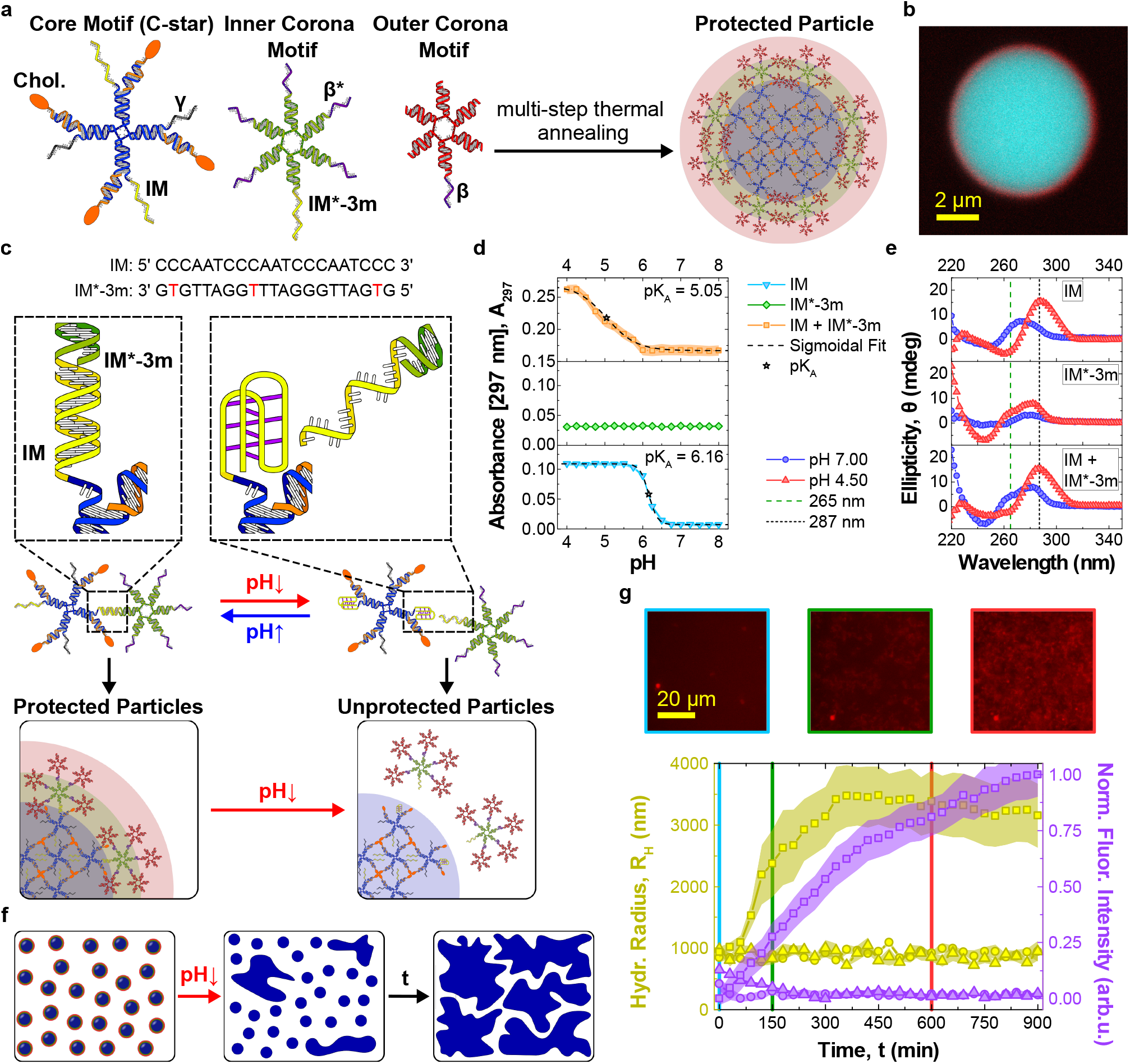
Core-shell DNA-based particles form a network in response to pH changes. **a**. Core-shell DNA particles assemble from cholesterolised (core motifs, CMs) and non-cholesterolised DNA nanostars (inner and outer corona motifs, ICM and OCM, respectively) [54]. CMs, composed of cholesterol-functionalised (orange) and non-functionalised strands (blue), form the hydrophobic particle core. ICMs and OCMs create a stabilising corona. CMs and ICMs bind through domains *IM* and *IM*^*∗*^*-3m*, while ICMs and OCMs attach through *β* − *β*\* overhangs. Domain *γ* can be used to fluorescently label CMs, by connecting to an Alexa Fluor 594-labelled duplex. When not used, *γ* is replaced by a poly-T sequence. Sequences of all strands used in this work and composition of all samples are outlined in Supplementary Tables 1 and 2, respectively. The relative thickness of core and shell region in the schematics is not in scale. **b**. Confocal micrograph of a large particle with distinguishable core-shell structure, assembled through a slow annealing protocol. Particles used in the remaining experiments had a much smaller size (200 nm – 1 *µ*m, see Methods). CMs are shown in cyan (fluorescein), OCMs in red (Alexa Fluor 647). **c**. *IM* and *IM*^*∗*^ − 3*m* domains are designed to cause the detachment of ICMs from CMs at low pH. *IM* is C-rich, able to form a non-canonical i-motif under acidic conditions [56], resulting in destabilisation of the duplex formed by *IM* and *IM*^*∗*^ − 3*m*. The duplex is rendered less stable by mismatches between the two sequences (red). **d**. pH-dependence of the UV absorbance at 297 nm, measured as proxy for i-motif formation [57, 58], for samples of *IM, IM*^*∗*^ −3*m* and *IM* + *IM*^*∗*^ −3*m* oligonucleotides (not linked to star motifs). Increase in *A*_297_ marks i-motif formation achieved at pH ∼6.16 for isolated *IM* and ∼5.05 when also *IM* ∗ −3*m* is present, with the difference ascribed to competition between duplex and i-motif formation. The transitional pH (pK_*A*_) values were calculated as the inflection points of sigmoidal fits. No response was observed in isolated *IM*^*∗*^ − 3*m*. The data are shown as mean ± standard error (shaded regions) of two experiments performed on two independently prepared samples, each consisting of three measurements. **e**. Circular dichroism (CD) spectra of the samples in panel **d**. Characteristic maxima at 287 nm and minima at 265 nm [57, 58] confirm i-motif formation in *IM* and *IM* + *IM*^*∗*^ − 3*m* samples. Data are averaged over three measurements. **f**. Schematic representation of pH-induced particle aggregation. pH decrease leads to i-motif formation, corona displacement, exposure of the sticky cholesterol-DNA cores, and ultimately particle aggregation. **g**. Bottom: particle aggregation after pH decrease tracked by measuring the hydrodynamic radius of growing aggregates with Differential Dynamic Microscopy (DDM, left axis) [59, 60] and the normalised epifluorescence intensity of accumulating CMs labeled with Alexa Fluor 594 (red, right axis). Triangles and squares represent responsive particles incubated at pH 7.0 (triangles) and 4.5 (squares). Circles indicate a control sample with nonresponsive particles at pH 4.5, where the *IM* and *IM*^*∗*^ − 3*m* domains have been replaced with non-responsive sequences. Data are plotted as mean ± standard error (shaded regions) of three (circles and triangles) or six (squares) measurements conducted on two independently prepared samples. Top: epifluorescence micrographs of responsive particles after pH decrease at different time points (*t* = 0, 150 and 600 min). See Supplementary Fig. 5 for additional micrographs.

Establishing this missing communication link requires the identification of a sufficiently general, cell-generated signal that can be coupled with the reconfiguration of the DNA nanostructures. Our model target – *E. coli* – similar to several other species, is well known to naturally acidify its microenvironment as a result of its sugar metabolism [55]. We thus identified pH as the ideal biogenous trigger, and proceeded to render the particles pH-responsive, so that the protective corona detaches from the sticky core at sufficiently low pH. This effect was obtained by engineering the DNA overhangs that link the CMs to the ICMs, and in particular by exploiting the pH-responsiveness of C-rich sequences. As highlighted in Fig. 2**c**, at neutral pH (∼ 7), the C-rich *IM* domain [56] on the CM hybridises to domain *IM* ^∗^ − 3*m* on the ICM, due to them being complementary but for three base mismatches. At lower pH, the duplex formed by *IM* and *IM* ^∗^ − 3*m* is destabilised by the formation of an intra-molecular, non-canonical i-motif in *IM* [56, 58, 64], thus leading to separation of the CM from the ICM.

We characterised the pH-induced i-motif-to-duplex transition by recording UV absorbance at 297 nm, known to increase following C-protonation and i-motif formation [57, 58]. Results, collated in Fig. 2**d**, show a transitional pH value (*pK*_*A*_) equal to 6.16 for samples that only contain *IM*, which is reduced to 5.05 when also *IM* ^∗^ − 3*m* is present, owing to duplex hybridisation counteracting i-motif formation. A similar *pK*_*A*_ was obtained with Dynamic Light Scattering measurements assessing the pH-dependent complexation of CMs and ICMs, as shown in Supplementary Fig. 4. i-motif formation was further confirmed by Circular dichroism (CD), as summarised in Fig. 2**e**. At pH 4.50, samples containing *IM*, both with and without *IM* ^∗^ − 3*m*, produced CD spectra with a maximum at 287 nm and minimum at 265 nm, both characteristic of the i-motif [57]. In traces recorded at pH 7.00, both extremes were shifted to lower wavelengths, as expected for dsDNA.

pH-induced corona displacement causes the exposure of the particles’ sticky cores, leading to their cholesterol-mediated aggregation and the formation of an extended amphiphilic DNA network, as shown in Fig. 2**f**. To assess network formation we used Differential Dynamic Microscopy (DDM) [59, 60], which allowed the monitoring of the time-evolution of the hydrodynamic radius of the particles, *R*_*H*_, after lowering the pH to 4.5. The data, collated in Fig. 2**g** (left axis, squares), show the expected increase in *R*_*H*_ from the initial value of ∼ 1 *µ*m, resulting from particle aggregation. Network formation is visible in epifluorescence micrographs collected using particles labelled with Alexa 594 (Fig. 2**g**, top). The mean (normalised) fluorescence intensity from the micrographs can further be used to track particle aggregation, exploiting progressive sedimentation of the aggregates that leads to an increase in the fluorescent signal close to the bottom of the imaging well. The fluorescence traces shown in Fig. 2**g** (right axis, squares), expectedly, follow a trend similar to the hydrodynamic radius. To confirm the specificity of the observed aggregation response, we conducted two control experiments, one in which pH responsive particles were kept at pH 7 (triangles) and another where non-responsive particles, obtained by replacing the C-rich motif with a sequence unable to form an i-motif, were exposed to pH 4.5 (circles). In both controls, no aggregation was noted from the *R*_*H*_ or fluorescence intensity data (Fig. 2**g**), confirming that the observed network formation indeed emerges from the designed pH-responsive pathway. The absence of non-triggered aggregation, also confirmed for medium compositions different from PBM9 (Supplementary Fig. 6), demonstrates the stability of the particles in bare media over experimentally relevant timescales. An extended set of bright field and epifluorescence micrographs, showing the behaviour of responsive/nonresponsive particles at neutral and acidic pH, is included in Supplementary Fig. 5.

### Bacteria immobilisation triggered by glucose metabolism in *E. coli*

Acidification of the extracellular milieu is a very common phenomenon, as a vast number of microorganisms produce acids as a consequence of fermentative processes. Particularly notable among them are bacteria belonging to the Bacillota and Pseudomonadota phyla that include human pathogens such as *Pseudomonas aeruginosa, Enterococcus faecalis, Salmonella enterica, Streptococcus pneumoniae, Yersinia pestis* and *Escherichia coli* [55]. Due to its high tractability in lab settings, we set out to demonstrate our synthetic netosis-like pathway in the presence of *E. coli*, which excretes acetate as a result of glucose metabolism thereby acidifying its growth medium, as sketched in Fig. 3**a**. Having designed responsive DNA-based particles capable of forming a sticky network upon exposure to acidic conditions, we proceed to determine whether *i)* the particles can be “activated” by glucose metabolism in *E. coli* and *ii)* the formed network can trap the cells, as required for the synthetic netosis-like response outlined in Fig. 1 (right).

**FIG. 3.**
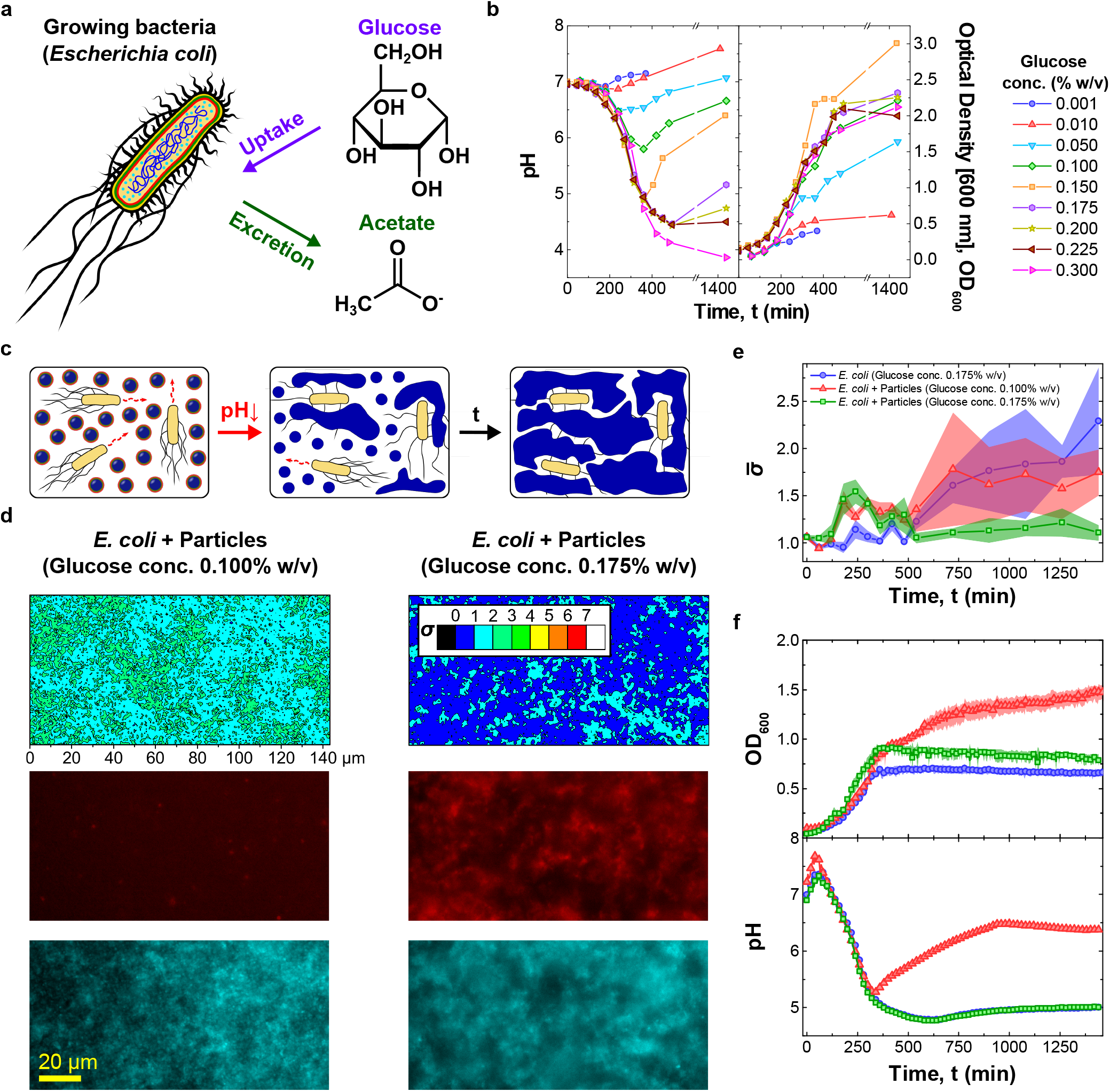
pH-responsive DNA particles trap *E. coli* when triggered by bacterial metabolism. **a**. Glucose metabolism in *E. coli* leads to acetate release and pH decrease [55]. **b**. (left) Bacteria-induced pH changes depend on glucose concentration in the medium, *ρ*_*G*_. For intermediate values (0.050% w/v ≤ *ρ*_*G*_ ≤ 0.200% w/v) the pH decreases and reaches a minimum before recovering. For *ρ*_*G*_ ≥ 0.225% w/v recovery is not observed. Glucose concentration influences bacterial growth, quantified through turbidity (OD) measurements at 600 nm (right). Culture yield increases with *ρ*_*G*_, is maximised at intermediate values, and decreases at higher glucose concentration, possibly due to excessive medium acidification. Data for *ρ*_*G*_ = 0.175, 0.200 and 0.225% w/v are shown as average of three independent repeats, the remaining points are from a single repeat. **c**. Diagram illustrating *E. coli* trapping by the synthetic DNA net. Bacterial metabolism reduces the pH, causing particle activation and the formation of a sticky network (Fig. 2) that embeds the cells. **d**. (top) *E. coli* immobilisation induced by DNA net formation as quantified through the motility parameter *σ*, extracted from bright field microscopy videos (see Methods). The two colourmaps are relative to samples containing *E. coli*, responsive particles and different *ρ*_*G*_ values, one insufficient (0.100% w/v, left) and the second sufficient (0.175% w/v, right) to reach the pH threshold for particle activation (5.05, see Fig. 2**d**). Bottom: epifluorescence micrographs corresponding to the *σ*-maps. Core motifs are labelled with Alexa Fluor 594 (red), while *E. coli* express EGFP (cyan). Smaller *σ*-values and co-localisation of DNA and bacteria observed in the sample with higher glucose concentration confirm the ability of cholesterol-DNA networks to bind and immobilise *E. coli. σ*-maps and images were collected at *t* = 1440 min after sample preparation. **e**. Time evolution of the frame averaged motility parameter 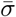 recorded for the samples in panel **d** and a control sample with *E. coli* and *ρ*_*G*_ = 0.175% w/v, but lacking DNA particles. The decrease in 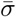 noted in the sample with responsive particles and higher glucose concentration confirms reduced *E. coli* motility following trapping. Data are shown as mean ± standard error of seven measurements conducted on three independent repeats. Associated *σ*-maps and epifluorescence micrographs are shown in Supplementary Fig. 11. **f**. Time-traces of OD (top) and medium pH (bottom) for the three samples in panel **e**. The pH was measured using the ratiometric pH probe FITC-dextran, added in solution (see Supplementary Figs 9 and 10, and Methods). For both OD and pH, data are shown as averages of three independent repeats. For the OD curves, standard errors are shown as shaded regions.

As demonstrated in Fig. 3**b** (left), the magnitude of the pH decrease in *E. coli* cultures can be modulated by changing the initial concentration (*ρ*_*G*_) of glucose in the medium. For all but the lowest tested glucose concentrations (0.001% and 0.01% w/v) a notable decrease in pH was observed, followed by a recovery for *ρ*_*G*_ ≤ 0.200% w/v due to re-uptake of acetate at later growth stages [55]. The magnitude of the pH drop correlates with glucose concentration, with values lower than 5, required for particle activation, reached when *ρ*_*G*_, 2 0.150% w/v. Figure 3**b** (right) explores the link between glucose concentration and cell growth, quantified through optical density (600 nm). The highest yield was observed for *ρ*_*G*_ = 0.150% w/v, with lower yields noted for both higher and lower *ρ*_*G*_ values. The substantially lower yield seen at high *ρ*_*G*_ (0.300% w/v) correlates with the lack of recovery in pH observed in Fig. 3**b** (left), hinting that excessive acidification may impact *E. coli* metabolism and prevent acetate re-uptake. In Supplementary Figs 7 and 8 we present OD and pH data collected for *E. coli* cultured in well plates, relevant for the synthetic netosis experiments discussed below (see Methods). Trends similar to those reported in Fig. 3**b** for flask cultures were noted, both in terms of growth curves and pH traces. A systematic difference was observed in the lowest pH reached for *ρ*_*G*_ = 0.300% w/v, which was higher in well plates compared to flask experiments (∼ 4.75 and ∼ 4, respectively). This deviation is likely due to the different method used to record pH, which relied on a physical pH probe in flasks and on a ratiometric fluorescent probe, FITC-dextran, in well plates. The response of the fluorescent probe was indeed found to saturate at pH ∼ 5 (Supplementary Fig. 9). We note that, to preserve the buffer conditions optimised for particle assembly and stability, bacteria were cultured in a newly developed medium dubbed PBM9. Medium composition, outlined in the Methods section, was selected to match ionic and buffering properties of PBS buffer and contain compounds needed to support bacterial growth, as in the M9 medium.

Having characterised the ability of *E. coli* to acidify their medium, we proceeded to test DNA-particle activation and consequent bacteria trapping. As sketched in Fig. 3**c**, the pH-responsive particles were deployed in *E. coli* samples with *ρ*_*G*_ = 0.175% w/v, sufficient to achieve the pH-level required for particle activation, and *ρ*_*G*_ = 0.100%, insufficient to reach the threshold (compare Fig. 2**d** (top) and Fig. 3**b** (left)). We note that the two *ρ*_*G*_ values result in comparable growth rates (Fig. 3**b**, right).

Epifluorescence images of both particles (red) and bacteria (cyan), collected after incubating the samples for *t* = 1440 minutes, are shown Fig. 3**d** (bottom). The snapshots readily demonstrate formation of the synthetic DNA net in the sample with higher glucose concentration, while no visible particle aggregation was seen for *ρ*_*G*_ = 0.100%. We can thus confirm that metabolism-induced acidification is capable of triggering particle aggregation, establishing the sought communication link between the cells and the synthetic DNA constructs. To assess whether the DNA net displays the desired ability to trap motile *E. coli*, a parameter *σ* was extracted from high-framerate bright-field microscopy videos of the samples, defined as the time-averaged standard deviation of the pixel-intensity computed over seven consecutive frames (see Methods). We have previously demonstrated that *σ* represents a good proxy for bacterial motility, taking larger values in samples of motile *E. coli* and lower values if the cells are immobilised [54]. A clear difference in *σ* between the sample with lower and higher *ρ*_*G*_ is notable from the colormaps in Fig. 3**d** (top), relative to the end-point of our experiments (*t* = 1440 minutes). The substantial reduction in bacterial motility observed for *ρ*_*G*_ = 0.175% confirms that the DNA-particles were able to trap *E. coli*, once activated by the bacteria themselves, hence confirming two-way particle-cell communication.

The relative homogeneity of *σ* across the fields of view allows us to track the time evolution of bacteria motility by monitoring the frame-averaged 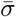, as summarised in Fig. 3**e** for the two samples discussed in Fig. 3**d**. A control *E. coli* sample with *ρ*_*G*_ = 0.175% w/v, but lacking DNA particles, is also included. After initial fluctuations, 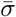 increases in the sample with *ρ*_*G*_ = 0:100% w/v and in the control sample, as a result of the increase in number of cells (compare Fig. 3**b**) and the lack of physical constraints limiting their motility. In contrast, in the sample containing particles, under conditions supporting their activation (*ρ*_*G*_ = 0.175% w/v), 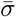 decreases at *t* = 510 min before plateauing at *t* = 720 min, as a consequence of DNA net formation and bacteria trapping. The onset of net formation correlates with the time-evolution of the pH reported in Fig. 3**f** (bottom), showing that the pH threshold required for particle activation (5.05) was reached prior to the time at which 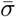 decreased. pH data were recorded with the fluorimetric probe FITC-dextran. See Supplementary Figs 9 and 10 for the calibration curve and raw fluorescence data, respectively. Both the pH trends and the OD growth curves shown in Fig. 3**f** (bottom and top, respectively) closely match data collected in samples lacking particles (Supplementary Fig. 8), indicating that particles have negligible effect on bacterial growth and metabolism under the tested conditions. Epifluorescence images and *σ* color-maps for the three samples discussed in Fig. 3**e**-**f** are shown in Supplementary Fig. 11 for selected time-points, enabling visual assessment of the difference in sample evolution resulting from net formation and bacteria trapping.

The modularity of our DNA nanodevices allowed us to introduce additional functionality in the bacteriarapping DNA particles. This is exemplified in Supplementary Figs 12 and 14, where we tested design variations f the particles that enable *in situ* fluorescent reporting of the bacteria-induced pH change, and consequent particle activation. In Supplementary Fig. 12, pH sensing was achieved by linking the ratiometric pH probe FITC-dextran to the core motifs. Observed trends were similar to the ones obtained with probes freely diffusing in bulk (Fig. 3**f** and Supplementary Fig. 10), but with the advantage that the labelled particles can probe pH in the local DNA-net microenvironment, in the vicinity of trapped bacteria. See also the calibration curve in Supplementary Fig. 13. In Supplementary Fig. 14, instead, we detected the pH-induced detachment of the corona motifs from the core motifs by placing a fluorophore and quencher on the *IM* and and *IM* ^∗^ − 3*m* domains, respectively. The biosensing capabilities of the material could be potentially further expanded by including agents that monitor alternative metabolic processes, *e*.*g*. Fe(III) respiration metabolism, as reported by Chen *et al*. [65].

### Bacterial growth inhibition induced by the synthetic netosis-like pathway

With the pH-responsive DNA particles we have successfully replicated two of the key responses of netosis, namely the detection of target cells and their immobilisation in a sticky DNA-cholesterol network. One last property is missing to fully mimic the biological response, and that is the antimicrobial action of the neutrophil extracellular traps causing cell death in the trapped pathogens. As observed above, however, the synthetic cholesterol-DNA net does not suppress bacterial growth under the tested conditions. Hence, to endow the system with antimicrobial properties, we introduced a second synthetic agent: a GUV loaded with the antibiotic ciprofloxacin, as sketched in Fig. 1**b**. The antibiotic was chosen due to its low minimum inhibitory concentration (MIC, Supplementary Fig. 15), and encapsulated within the GUVs at 200 × this value (3.2 *µ*g mL^−1^). The designed netosis-like response of the resulting three-agent artificial consortium is graphically outlined in Fig. 4**a**: after *E. coli* -induced medium acidification, the unprotected particles are expected to bind to the GUV and either disrupt or permeabilise them, causing the release of the drug that would, in turn, hinder the growth of trapped bacteria. The ability of unprotected cholesterol-DNA particles to render GUVs permeable has been tested in our previous contribution, using a fluorescent probe [54].

**FIG. 4.**
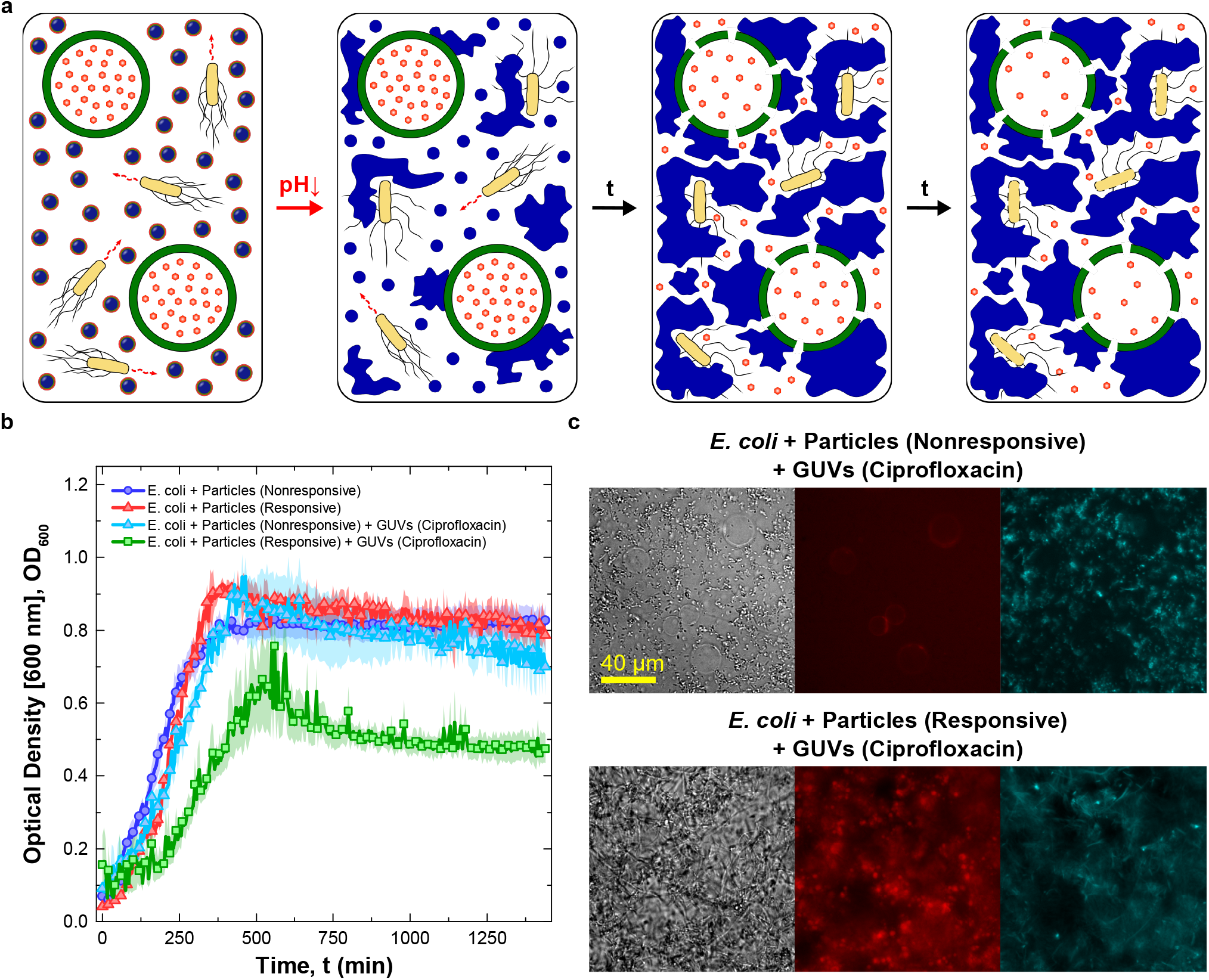
Synthetic cell signalling network produces netosis-like response. **a**. Diagram illustrating the mechanism of action of the synthetic signalling network producing a netosis-like response. Medium acidification caused by *E. coli* glucose metabolism activates the DNA-particles. The particles form a sticky DNA-cholesterol network that, simultaneously, traps the bacteria and permeabilises Giant Unilamellar Vesicles (GUVs) loaded with antibiotic ciprofloxacin. The released antibiotic hinders bacterial growth. **b**. Antimicrobial response as determined *via* turbidity measurements (OD at 600 nm). Data are shown for four samples as the mean of three independent repeats *±* standard errors (shaded regions). One sample includes *E. coli*, antibiotic-loaded GUVs, responsive particles, and sufficient glucose to achieve their activation (*ρ*_*G*_ = 0.175% w/v), and is thus capable of supporting the cascade of reactions producing the sought netosis-like response (green squares). The other tree samples are controls missing one or more key components. Two control samples lack antibiotic-loaded GUVs, and feature either non-responsive (blue circles) or responsive (red triangles) DNA particles. The third control sample contains antibiotic-loaded GUVs but uses non-responsive particles. While all control samples show similar OD curves, growth is delayed and suppressed in the system capable of sustaining the designed cascade of reactions. **c**. Bright field and epifluorescence micrographs of samples containing *E. coli*, nonresponsive (top) or responsive (bottom) particles and ciprofloxacin-loaded GUVs, recorded at *t* = 1440 min after sample preparation. Aggregation of DNA and the lack of GUVs in the sample with responsive particles indicates the successful rupture of vesicles following DNA-net formation. The released antibiotic hinders *E. coli* division, causing cell elongation. Particles (core motifs) are shown in red (Alexa Fluor 594), *E. coli* in cyan (EGFP). See Supplementary Fig. 16 for additional bright field and epifluorescence micrographs recorded at *t* = 0 and 1440 min.

To evaluate the antimicrobial action of the drug-loaded GUVs, and thus validate the netosis-like response of our synthetic consortium, we conducted an assay in which GUVs and pH-responsive particles were deployed in an *E. coli* sample along with 0.175% w/v glucose, sufficient to induce particle activation thorough medium acidification (Fig. 3). The resulting OD growth curve is shown in Fig. 4**b** (green squares), and compared with data from three distinct control experiments where we selectively eliminated components and functionalities required for the antimicrobial response. These include samples in which GUVs were not present and particles were either non-responsive (blue circles) or pH-responsive (red triangles), or where the GUVs were present along with non-responsive particles (cyan triangles). All control ex-periments produced similar growth curves, providing a baseline and confirming that neither the particles (responsive or non responsive), nor undisturbed antibiotic-loaded GUVs, influence growth. The sample sustaining the complete netosis-like pathway, instead, produced a visibly different response. While growth is similar to the controls in the initial two hours, once acidification and consequent particle activation occur, growth slows down leading to a maximum OD lower compared to the controls, followed by a noticeable decrease in OD at later times. A visual insight in the response of the systems can be gained from the bright field and epifluorescence images in Fig. 4**c**, collected at the end of the experiments (1440 minutes from sample preparation). Here, the control sample featuring GUVs and non-responsive particles is compared with the complete netosis-like system. In the control, no DNA net was formed (red epifluorescence channel) and the GUVs clearly retained their structural integrity (bright field) despite weak interactions with the non-activated particles (red channel). Bacteria also retained their physiological rod-like shape (bright field and cyan epifluorescence channel). Instead, in the sample capable of sustaining the full cascade of reactions leading to netosis, GUVs were no longer visible, having been disrupted by the formed DNA net. The consequent release of antibiotic had a clear effect the ability of *E. coli* to divide, as the cells acquired a visible filamentous morphology. The microscopy data, along with additional images provided in Supplementary Fig. 16, confirm the ability of our synthetic consortium to sustain the designed responses leading to the sought netosis-like mechanism.

## DISCUSSION

In this work we reported the bottom-up construction of a synthetic signalling network capable of imitating a complex immune response, namely that of netosis. Through this process, neutrophils detect, trap, and kill pathogens by secreting a DNA-based extracellular trap. Our artificial netosis pathway relies on the action of DNA-based synthetic particles, which upon detecting bacteria-induced medium acidification, form a sticky network reminiscent of the biological trap, capturing the cells. Simultaneously, the particles cause antibiotic to be released from liposomes, hindering growth and division in the trapped cells.

The netosis-like behaviour emerges from the coordinated activation of multiple biomimetic functions, including biosensing, morphological adaptation and communication, exemplifying how advanced life-like responses can be implemented with a relatively small number of molecular components, exploiting the modularity and programmability of nucleic-acid nanotechnology. Inspired by our solution, similar synthetic pathways could be implemented to program responses in the bacterial communities different from their death, for instance by loading the liposomes with inducers triggering protein synthesis [50], exploiting the cell-trapping functionality to engineer multi-species microbial consortia [66], or scaffolding synthetic biofilms, valuable in biomaterial synthesis [67] and bioremediation [68, 69]. The resulting “living biomaterials” [70], may also feature rheological properties which are dependent on the activity of the trapped cells, deserving future investigation [71]. Matrix rheology could, in turn, influence cell growth [72], enabling the design of adaptable materials reliant on biomechanical feedback loops.

Besides pushing the boundaries of bottom-up synthetic biology, our implementation could underpin disruptive antimicrobial solutions that, like our natural immunity, benefit from the combination of physical (cell immobilisation) and chemical (antibiotics) modes of action. To this end, activation of the synthetic net could be rendered conditional to pathogen-associated stimuli other than pH, such as the presence of cell-surface antigens [73] or soluble biomarkers [74], exploiting base-paring, aptamer technology or functionalisation with chemical or biological ligands. Solutions could also be devised to integrate the cargo-loaded liposomes and DNA-particles in a single agent, exploiting well-established strategies to link DNA devices to lipid membranes [75, 76]. The resulting “synthetic immune cell” could be valuable beyond antimicrobials, and be adapted to tackle, *e*.*g*., inflammation or cancer by adjusting the therapeutic payload and trigger stimulus, building towards the ambitious goal of creating effective and robust synthetic cellular therapies [42].

## METHODS

### Design and handling of DNA strands

DNA nanostructures were designed in NUPACK [77]. A constraint forbidding more than three consecutive C or G bases was imposed to inhibit formation of unwanted secondary structures. The i-motif forming domain and its complement were excluded from this constraint. All sequences are provided in Supplementary Table 1. Core Strand 1, carrying the internal Rhodamine 6G modification was purchased from Eurogentec. All remaining strands were bought from Integrated DNA Technologies (IDT). The functionalised and non-functionalised oligonucleotides were purified by the supplier using high-performance liquid chromatography (HPLC) and standard desalting, respectively. Once delivered, the dehydrated DNA was reconstituted in phosphate-buffered saline (PBS, 137 mM NaCl, 2.7 mM KCl, 8 mM Na_2_HPO_4_, 2 mM KH_2_PO_4_, pH 7.4, Invitrogen, Thermo Fisher Scientific). To remove any large particulate contaminants the solutions of non-functionalised oligonucleotides were syringe-filtered with polyethersulfone filters (0.22 *µ*m pore size, Millex). The concentration of reconstituted DNA samples was determined by measuring the absorbance at 260 nm on a Thermo Scientific Nanodrop 2000 UV–Vis spectrophotometer. The extinction coefficients were provided by the supplier (IDT) or calculated using the OligoAnalyzer online tool from IDT. All stock solutions were stored at -20^°^C.

### Preparation of non-functionalised DNA structures

Samples used to probe the folding efficiency, correct binding, thermal stability and pH-responsiveness of individual nanostructures were prepared from non-functionalised strands to prevent aggregation mediated by hydrophobic moieties. To achieve this, all the cholesterol-functionalised oligonucleotides were replaced with non-modified strands of identical sequence.

Concentrated solutions of DNA strands in PBS were prepared by stoichiometrically mixing all the required oligonucleotides in 200 *µ*L DNase free Eppendorf tubes. Note that, at this stage, the concentration of DNA strands was set to be 2 × or 2.15 × higher as compared to the final values given in Supplementary Table 2, to allow introduction of the M9 component of the PBM9 medium (see relevant Methods section below) and pH adjustment. Prepared mixtures were placed in a Techne TC-512 thermal cycler, heated up to 95^°^C, held at this temperature for 5 min, and then cooled down to 20^°^C at the rate of − 0.05^°^C min^−1^ to enable nanostructure assembly.

Samples for agarose gel electrophoresis (AGE) and UV melting curves of non-cholesterolised DNA structures (60 *µ*L and 600 *µ*L, 2 × DNA concentration) were transferred into 1.5 mL DNase free Eppendorf tubes and mixed in equal volumes with a solution containing PBS buffer, 37.4 mM NH_4_Cl, 0.2 mM CaCl_2_, 4 mM MgSO_4_, 0.4% w/v casamino acids (CAAs) and 0.35% w/v glucose (pH 6.54), termed PB(2 ×)M9, to obtain 1 × DNA concentration in PBM9 medium (pH 7.00).

For the preparation of samples for dynamic light scattering (DLS), UV-vis absorbance, and circular dichroism (CD) of non-functionalised DNA nanostructures, 700 *µ*L of 2.15 × concentrated solutions of assembled DNA nanostructures were placed in 50 ml centrifuge tubes (Greiner Bio-One) and supplemented with 700 *µ*L of PB(2 ×)M9. Afterwards, their pH was tuned with hydrochloric acid (HCl, 1 M or 6 M, Sigma-Aldrich) or potassium hydroxide (KOH, 1 M or 5 M, Sigma-Aldrich) solutions using a bench-top pH meter (pH 210, Hanna Instruments) equipped with a double junction pH electrode (9110DJWP, Orion). Finally, their volume was adjusted to 1500 *µ*L by adding milli-Q water.

All the prepared samples were stored at 4^°^C and used within 24 h.

### Preparation of pH-responsive particles

DNA strands were mixed in stoichiometric ratio in PBS to the final concentration of core strands equal to 1 *µ*M (concentrations of all the oligonucleotides are listed in Supplementary Table 2), in 500 *µ*L DNase free Eppendorf tubes. The overall sample volume was set to 60 *µ*L or 200 *µ*L, depending of the annealing protocol used.

Small volumes (60 *µ*L) of the samples used to prepare large particles (Fig. 1**b**), were loaded into borosilicate glass capillaries (inner section of 4 mm × 0.4 mm, CM Scientific). The capillaries were previously cleaned by sonicating for 15 min once in a 2% Hellmanex III water solution (Hellma Analytics) and then twice in milli-Q water. Both ends of the capillaries were capped with mineral oil, and sealed shut and attached to microscope slides (Menzel Gläser, 24 mm × 60 mm, No. 1) with fast-drying epoxy glue (Araldite). Sealed capillaries were then placed in a custom-made Peltier-controlled water bath and heated up to 90^°^C. After incubating for 30 min, the samples were cooled down to 63^°^C at the rate of − 0.01^°^C min^−1^ and finally brought down to 20^°^C at −0.1^°^C min^−1^.

pH-responsive particles of controlled size were prepared in a custom-made Peltier-controlled aluminium chamber allowing rapid temperature changes. 200 *µ*L volumes of the DNA strand mixtures were pipetted into ExtraSil (ES) quartz cuvettes (350 *µ*L, 1 mm path length, 10 mm inside width, Aireka Cells) using gel-loading pipette tips (1-200 *µ*L, 0.5 mm thick round end, RNase/DNase free, Corning). Loaded cuvettes were placed in the aluminium chamber, heated up to 90^°^C and incubated for 15 min. The samples were then cooled down to 65^°^C at −1^°^C s^−1^ and held at this temperature for a growth time *t*_*g*_ = 15, 600 or 900 s depending on the desired particle size. Subsequently, the temperature was decreased to 35^°^C at − 1^°^C s^−1^, held at this value for 15 min to facilitate corona formation and then further decreased to 20^°^C at the same rate. The particles can be melted and re-annealed multiple times, and we expect that one should be able to re-assemble the particles after pH-induced network formation, as long as the pH is brought back to the initial value and the DNA density is not substantially altered.

### Agarose gel electrophoresis of non-functionalised DNA motifs

AGE was used to asses the correct folding and binding of non-functionalised DNA motifs as shown in Supplementary Fig. 1.

Samples of annealed DNA nanostructures, prepared as described above, were first mixed with 4 *µ*L of loading dye (Thermo Fisher Scientific) and then diluted in Trisborate-EDTA (TBE) buffer (pH 8.3, 89 mM Tris-borate, 2 mM EDTA, Sigma-Aldrich) to the final DNA concen-tration of 75 ng mL^−1^.

Agarose gels were prepared at 1.5% w/v agarose (Sigma-Aldrich) in TBE buffer supplemented with SYBR safe DNA gel stain (Invitrogen, Thermo Fisher Scientific). Wells were filled with small volumes (10 *µ*L) of diluted samples, equivalent to 750 ng of DNA. The two outermost wells carried a 100 bp DNA reference ladder (Thermo Fisher Scientific). Gels were run for 120 min at 75 V (3 V cm^−1^), and then imaged using a GelDoc-It imaging system.

The collected images were analysed through a tailormade Matlab script to obtain the integrated line intensity profiles.

### Dynamic light scattering of non-cholesterolised nanostar structures

DLS was used to both validate the successful assembly of non-cholesterolised DNA nanostructures and examine their stability at acidic pH (see Supplementary Fig. 2). First, an ultra low volume quartz cuvette (ZEN2112, Malvern) was loaded with 120 *µ*L of annealed sample in PBM9 medium with preadjusted pH that was set to either 7.0 or 4.5. Afterwards, three measurements, each consisting of fifteen data runs, were taken at room temperature with a Malvern Zetasizer Nano ZSP analyzer (scattering angle fixed at 173^°^, 633 nm He–Ne laser). For each of the visible bands, the hydrodynamic diame-ter *D*_*H*_ was determined by fitting the experimental data to a lognormal distribution function and then calculating the maximum of the fit, using a custom Matlab script.

### UV melting curves of non-functionalised DNA structures

UV-Vis spectrophotometry was used to asses the separation of melting temperatures *T*_*M*_ of core-forming motifs and corona-binding domains (Supplementary Fig. 3), required by the multi-step thermal annealing process used to form core-shell particles.

Large volumes (1200 *µ*L) of samples containing non-functionalised DNA structures, prepared according to the protocol reported above, but excluding the annealing step, were loaded into quartz cuvettes. To prevent evaporation, 300 *µ*L of mineral oil were carefully pipetted on top of each sample before sealing the cuvettes with Parafilm-wrapped polytetrafluoroethylene (PTFE) stoppers.

Samples were first cooled down from 95^°^C to 20^°^C and then heated back up to 95^°^C at the rate of ± 0.02^°^C min^−1^ while their absorbance at 260 nm was measured on a temperature-controlled Varian Cary 50 UV-Vis spectrophotometer.

*T*_*M*_ was determined with a custom Matlab script for both the cooling and heating ramps by fitting the upper and lower plateaus with straight lines and then computing the intersection between their median and the experimental data *via* interpolation [78].

### Confocal microscopy of large core-shell particles

Confocal microscopy images of large particles (Fig. 2**b**) were obtained with a Leica TCS SP5 laser scanning confocal microscope equipped with a HC PL APO CORR CS 40 × /0.85 dry objective (Leica).

Aggregates, prepared according to the procedure described above, were extracted from the capillary by cutting open its ends with a diamond scribe, placing it in 1.5 mL DNase free Eppendorf tube containing 60 *µ*L of PBS buffer, and spinning down with a tabletop centrifuge for ∼ 30 s. The sample extracted sample was then washed twice to remove the surplus corona motifs by first diluting in PBS to the total volume of 120 *µ*L, followed by centrifugation at 420 *g* for 30 min and supernatant replacement with fresh PBS buffer (90 *µ*L). Washed aggregates were pipetted into silicone rubber incubation chambers (6.5 mm × 6.5 mm × 3.5 mm, Grace Biolabs FlexWells) and sealed with DNase free tape (Grace Biolabs FlexWell Seal Strips) to prevent evaporation.

For imaging, an Ar-ion laser line at 488 nm and an HeNe line at 633 nm were used for excitation of fluorescein (core motifs) and Alexa Fluor 647 (outer corona motifs) dyes, respectively.

### Assessment of i-motif formation with UV absorbance

UV-Vis spectrophotometry was used to characterise i-motif formation (Fig. 2**d**).

Annealed oligonucleotide samples (1200 *µ*L) in PBM9 medium with pre-adjusted pH were loaded into quartz cuvettes, and their absorbance at 297 nm [57, 58] was measured at room temperature using the aforementioned UV-Vis spectrophotometer. The tested pH ranged from 8 to 4 with steps of 0.25 points.

The transitional pH values (pK_*A*_) were calculated with a tailor-made Matlab script as the inflection point of sigmoidal fits to the experimental data.

### Assessment of i-motif formation with DLS

DLS measurements, performed on the Malvern Zetasizer Nano ZSP setup, were used to confirm the accurateness of the transitional pH value extracted from UV response curves (Fig. 2**d**). The recorded pH-response curve is shown in Supplementary Fig. 4.

For these experiments, 120 *µ*L of the samples containing core and inner corona motifs in PBM9 medium with pre-adjusted pH were pipetted into an ultra-low volume quartz cuvette. Three measurements, each consisting of fifteen data runs, were then taken at room temperature. The tested pH ranged from 8 to 4 with steps of 0.25 points.

The plotted *D*_*H*_ values were extracted from raw intensity profiles using the previously mentioned Matlab script.

### Assessment of i-motif formation with circular dichroism

CD measurements, performed on a JASCO J-810 CD spectrophotometer, were used to further validate the imotif formation upon pH decrease (Fig. 2**e**).

First, 200 *µ*L of highly concentrated (20 *µ*M) samples in PBM9 medium with pH adjusted to 7.00 or 4.50 were pipetted into a 1 mm path-length, stoppered quartz cuvette. CD spectra were then acquired at room temperature in the spectral range of 210-350 nm.

### pH-triggered particle aggregation assay with epifluorescence and differential dynamic microscopy

Differential dynamic microscopy (DDM) and epifluorescence microscopy were used to asses the pHresponsiveness of the DNA particles (Fig. 2**g** and Supplementary Fig. 5).

Responsive (two samples) and nonresponsive (one sample) particles prepared in PBS buffer with the thermal annealing protocol described above (*t*_*g*_ = 900 s) were transferred into silicon incubation chambers and supplemented with PB(2×)M9 at 1:1 ratio to the final volume of 120 *µ*L. The pH of two of the samples was then adjusted with HCl and KOH to 4.50 before sealing the wells with DNase free tape and imaging for 960 min at 30 min intervals. The remaining sample with responsive particles served as a control. Both high frame-rate bright field videos (150 fps, 8 s) and epifluorescence micrographs were recorded with a fully motorised Nikon Eclipse Ti-E inverted epifluorescence microscope, equipped with CFI Plan Apochromat *λ* 40 × /0.95 NA dry objective (Nikon), Grasshopper3 GS3-U3-23S6M camera (Point Gray Research) and Perfect Focusing System (Nikon). For the epifluorescence imaging of core motifs labelled with Alexa Fluor 594, red LED illumination and a Texas Red filter cube (Semrock) were used.

Bright field videos were analysed with DDM [59, 60] using a custom script written in Matlab, as described in ref. [54].

Briefly, an image structure function, Δ*I*(*q, τ*), was obtained as

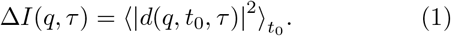

In Eq. 1, *q* is the wave-vector in Fourier space and *d*(*q, t*_0_, *τ*) is the 2D spatial Fourier transform of *d*(*r, t*_0_, *τ*) = *I*(*r, t*_0_ + *τ*) − *I*(*r, t*_0_), where *I*(*r, t*_0_) and *I*(*r, t*_0_ + *τ*) are two video frames collected at times *t*_0_ and *t*_0_ + *τ*. Δ*I*(*q, τ*) was then fitted to [59]

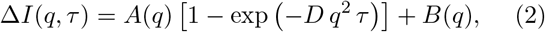

to obtain the Brownian diffusion coefficient *D. A*(*q*) and *B*(*q*) are functions dependent on the static scattering properties of the sample, instruments optics and camera noise.

The computed *D* was used to calculate the hydrodynamic radius *R*_*H*_ from the Stokes-Einstein equation. Note that at late times, once aggregates become too large to diffuse and/or start percolating, the extracted *R*_*H*_ looses its physical meaning. Despite this, all experimental data were fitted to Eq. 2 for consistency.

Analysis of epifluorescence micrographs was performed with a tailor-made Matlab script by calculating the sum of the background-subtracted pixel intensity within the observed field of view.

### Particle stability in bacterial growth media

DDM was used to evaluate the stability of DNA particles after deployment in several types of commonly used bacterial growth media (Supplementary Fig. 6).

Small volumes (60 *µ*L) of nonresponsive particles prepared in PBS buffer using the thermal annealing protocol described above (*t*_*g*_ = 600 s) were transferred into three silicon incubation chambers. The wells were then sealed with DNase free tape and imaged by recording three high frame-rate bright field videos (150 fps, 8 s) per sample with the previously described Nikon Eclipse Ti-E inverted epifluorescence microscope.

After 5 min, the chambers were supplemented with M9 medium (12.8 g L^−1^ Na_2_HPO_4_ 7H_2_O, 3 g L^−1^ KH_2_PO_4_, 0.5 g L^−1^ NaCl, 1 g L^−1^ NH_4_Cl, 11 mg L^−1^ CaCl_2_, 240 mg L^−1^ MgSO_4_ and 0.4% w/v Glucose), M63 medium (13.6 g L^−1^ KH_2_PO_4_, 0.5 mg L^−1^ FeSO_4_ 7H_2_O, 0.5 g L^−1^ MgSO_4_ 7H_2_O, 1.27 mg L^−1^ Thiamine, 2.64 g L^−1^ (NH_4_)_2_SO_4_ and 0.5% w/v Glucose) or M63 medium containing casamino acids (CAAs, 0.4% w/v) at 1:1 ratio to the final volume of 120 *µ*L. After this, the wells were resealed and the samples were imaged by acquiring three high frame-rate bright field videos per chamber at times *t* = 60, 120, 180, 300 and 1200 min from the growth medium addition.

The acquired data was analysed using a custom DDM analysis script written in Matlab, as described in the previous section.

### Bacterial strains and growth conditions

All the experiments involving bacteria were carried out with the highly motile MG1655 *Escherichia coli* strain. For fluorescence imaging (Figs 3**d**, 4**c** and Supplementary Figs 10, 16), the strain was transformed with the pWR20-EGFP plasmid that enables the constitutive expression of the enhanced green fluorescent protein (EGFP) in the cytoplasm and kanamycin resistance [79, 80].

*E. coli* cells were grown from single colonies in 50 ml conical glass flask filled with 5 mL of LB medium (10 g L^−1^ Tryptone, 5 g L^−1^ Yeast extract, 0.5 g L^−1^ NaCl) in an overnight incubation carried out at 37^°^C, with continuous shaking at 220 rpm. A small volume (50 *µ*L) of the culture in LB was then transferred into a glass flask containing 50 *µ*L of M63 medium and the cells were allowed to grow to an optical 0.2-0.3 OD using the same incubation conditions. Before the experiments, bacteria were pelleted through centrifugation at 8000 *g* for 2 min, washed twice with fresh buffer (PBS or PB(2 ×)M9) and concentrated to obtain a final OD of 0.11 after dilution by mixing all the components required to perform specific experiments. Note that to guarantee the expression of EGFP in all *E. coli*, both LB and M63 media were supplemented with kanamycin.

### PBM9 medium

The deployment of core-shell DNA particles in bacterial cultures yields solutions that are partly made up of PBS buffer used in their production. To robustly grow bacteria in a medium that matches the ionic and buffering characteristics of PBS buffer, we devised PBM9 (137 mM NaCl, 2.7 mM KCl, 8 mM Na_2_HPO_4_, 2 mM KH_2_PO_4_, 18.7 mM NH_4_Cl, 0.1 mM CaCl_2_, 2 mM MgSO_4_, 0.2% w/v CAAs, various concentrations of glucose, pH adjusted to 7.00 with HCl and KOH) which is based on the common M9 medium, with the buffering and ionic components of PBS.

The medium was used in all the experiments included in this study that involve *E. coli* and/or DNA particles/nanostructures.

### Bacterial growth and pH changes in PBM9 medium

*OD* measurements, performed on the aforementioned UV-Vis spectrophotometer, were used to characterise the bacterial growth in PBM9 medium (Fig. 3**b**). The accompanying pH changes were tracked using a benchtop pH meter. 250 *µ*L of *E. coli* cells in LB (overnight growth) were loaded into 250 ml conical glass flasks with 24.75 mL of PBM9 medium (various glucose concentrations) and then incubated at 37^°^C with continuous shaking at 220 rpm. Every 45 minutes, two volumes of 1 ml each were transferred into 1.5 ml Eppendorf tubes. The first sample was pipetted into a disposable cuvette (polystyrene, BRAND) and used to measure *OD*. After the measurement, the aliquot was transferred back into the flask. The second sample was filtered using a 0.22 *µ*m polyethersulfone filter into 50 ml centrifuge tube to remove bacteria cells. The resultant solution was used to measure pH. The procedure described above was repeated eleven times at *t* = 45, 90, 135, 180, 225, 270, 315, 360, 405, 450 and 495 min, after which the samples were left incubating for additional 960 min with the final measurement performed at the end of incubation cycle.

### Plate reader-based assay of pH and optical density changes accompanying the bacterial growth

Turbidity and fluorescence measurements, performed on a FLUOstar Omega plate reader, were used to determine the optimal glucose concentration enabling bacteria-induced particle activation in conditions used for further experiments (Supplementary Figs 7, 8).

*E. coli* cells were initially grown and washed as described above, then resuspended in PBS and loaded into a DNase free, flat-bottom 96-well plate (CELLSTAR Black 96 Well Cell Culture Microplates, Greiner Bio-One). Each well was then topped with 120 *µ*L of PB(2×)M9 (various glucose concentration) and 10 *µ*L of fluorescein (FITC)dextran (3 kDa, anionic, Sigma-Aldrich) solution in PBS to a total volume of 240 *µ*L, resulting in the final FITC-dextran concentration of 50 *µ*M and OD of 0.11. OD and fluorescence intensity at 520 nm upon excitation at 400 nm (*I*_400_) and 485 nm (*I*_485_) were measured in the plate reader for 2160 min at 8 min intervals while shaking at 700 rpm in double-orbital mode and keeping the temperature at 37^°^C.

The pH was extracted from the intensity ratio *I*_485_*/I*_400_ [81] using a calibration curve (Supplementary Fig. 9), The calibration curve was acquired by first preparing 1 mL solutions of 50 *µ*M FITC-dextran in PBM9 (various pH), loading 240 *µ*L of each sample into a 96-well plate, and then measuring *I*_400_ and *I*_485_ on the plate reader at 37^°^C. The pH ranged from 8 to 4 at 0.25 intervals, and was adjusted with HCl and KOH.

The collected data were processed with a tailor-made Matlab script by fitting with a sigmoid function.

### Bacteria immobilisation assay

For experiments on bacteria-induced particle aggregation and subsequent *E. coli* immobilisation (Figs 3**d, e** and Supplementary Fig. 11), three wells in a 96-well plate were filled with 10 *µ*L of highly concentrated *E. coli* solution in PBS (*OD* = 0.66 after 4 × dilution), prepared as described above, 10 *µ*L of PBS and 120 *µ*L of PB(2×)M9 (glucose concentration set at 0.100% w/v or 0.175% w/v after dilution at 1:1 ratio). Two of the wells with different glucose content were topped with 100 *µ*L of the PBS solution containing pH-responsive particles (*t*_*g*_ = 15 s), while the third well was filled with an identical volume of PBS, and used as a control. Afterwards, samples were incubated in the previously introduced plate reader for *t* = 0, 60, 120, 180, 240, 300, 360, 420, 480, 540, 720, 900, 1080, 1260 and 1440 min at 37^°^C, with shaking at 500 rpm for 30 s every 5 min. Note that a distinct set of samples was used for each time point.

All samples were imaged with the aforementioned bright field and epifluorescence microscopy setup by taking a set of seven high frame-rate videos (150 fps, 8 s) and epifluorescence micrographs. Green LED illumination/Texas Red filter (Semrock) and blue LED/GFP filter (Semrock) were used to record signals from core motifs (Alexa Fluor 594) and *E. coli* (EGFP), respectively.

The parameter *σ*, introduced in ref. [54] and used here to gauge bacterial motility (Fig. 3**d** and Supplementary Fig. 11), was calculated with a custom Matlab script from bright field videos as 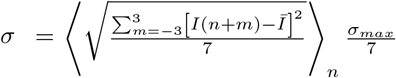, where *I*(*n*) represents the *n*-th frame of video, 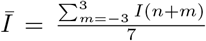 dicates a rolling average over the entire video, and *σ*_*max*_ is the highest intensity standard deviation value recorded in all the videos. Parameter 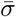 shown in Fig. 3**e** was computed by averaging *σ* over all the pixels in a frame.

Bacterial growth and pH changes occurring throughout the assay (Fig. 3**f** and Supplementary Fig. 10) were monitored on the plate reader with analogous samples containing non-fluorescent particles and *E*.*coli* lacking the pWR20-EGFP plasmid. To enable pH tracking, 10 *µ*L of PBS were replaced with an identical volume of 1.2 mM FITC-dextran solution in PBS. Prepared samples were incubated for 1440 min at 37^°^C with shaking at 500 rpm for 30 s every 5 min. *OD, I*_485_ and *I*_400_ were measured at 5 min intervals over time. The pH was then calculated from *I*_485_*/I*_400_ using the calibration curve shown in Supplementary Fig. 9.

### Bacteria detection with a ratiometric pH probe

For experiments used to sense pH changes with fluorophore-labelled particles, (Supplementary Fig. 12), two wells in a 96-well plate were filled with 20 *µ*L of PBS solution containing *E. coli* cells (*OD* = 0.66 before concentrating the cells 2×). Both chambers were then topped with 100 *µ*L of responsive particles (*t*_*g*_ = 15 s) in PBS, which were formed using fluorescein-labelled core motifs, and 120 *µ*L of PB(2×)M9 medium with different glucose concentrations (0.100% w/v or 0.175% w/v in the final solution). Two additional wells were loaded with 240 *µ*L of equimolar particle solutions in PBM9. The pH of one of them was adjusted to 4.50 with HCl and KOH. Fluorescence intensities *I*_400_ and *I*_485_ were automatically measured in the aforementioned plate reader for 1440 min, at 5 min intervals, temperature set to 37^°^C, with shaking at 500 rpm for 30 s every 5 min.

The recorded fluorescence intensity ratios *I*_485_*/I*_400_ were used to calculate pH using calibration curve recorded for fluroescein-dextran probes (Supplementary Fig. 9). The use of this calibration curve is justified by the control experiment in Supplementary Fig. 13.

### Bacteria sensing with a fluorophore-quencher pair

Fluorescence measurements used to sense bacteria-induced pH changes with particles modified with the fluorophore-quencher pair (Supplementary Fig. 14) were carried out on the plate reader described above. Rhodamine 6G (Rh6G) and Black Hole Quencher 1 (BHQ1) were used as fluorophore and quencher, respectively.

First, small volumes (10 *µ*L) of *E. coli* solution in PBS (4 × dilution results in OD of 0.66), prepared as in the protocol described above, were loaded into four wells of a 96-well plate. Two of the chambers were then filled with 100 *µ*L of responsive particles (*t*_*g*_ = 15 s, Rh6G and BHQ1) in PBS, 10 *µ*L of *IM*-3m* (BHQ1) strand in PBS, and 120 *µ*L of PB(2×)M9 medium. The BHQ1-modified strand was included to block any remaining sites with the Rh6G dye located within the particles’ cores, that could not be accessed and quenched by inner corona motifs during particle assembly. The two remaining wells were topped with 110 *µ*L of PBS buffer and 120 *µ*L of PB(2×)M9 medium. The overall glucose concentration was set to 0.100% w/v or 0.175% w/v and varied between samples of the same type. In the next step, a control sample was prepared in a separate well by mixing 100 *µ*L of PBS solution containing responsive particles (*t*_*g*_ = 15 s, Rh6G) with 20 *µ*L of PBS buffer and 120 *µ*L of PB(2 ×)M9 medium. All the samples were incubated in the plate reader for 1440 min at 37^°^C, with shaking at 500 rpm for 30 s every 5 min. Fluorescence intensity at 530 nm upon excitation at 485 nm was monitored over time every 5 min.

The final curves representing the samples that contain both *E. coli* and particles in PBM9 medium with distinct glucose concentration, shown in Supplementary Fig. 14, were obtained by subtracting the signal recorded from analogous samples lacking the particles. This operation was performed to eliminate contributions from *E. coli* autofluorescence, which can be detected at this wavelength [82].

### MIC estimation assay

The minimum inhibitory concentration (MIC) of the antibiotic ciprofloxacin was measured using the broth microdilution technique [83] on the aforementioned plate reader (Supplementary Fig. 15).

Cells were grown in LB at 37^°^C overnight with shaking at 220 rpm, and inoculated at a 1:100 dilution into a 96-well plate containing various concentrations of ciprofloxacin in PBM9, for a final volume of 200 *µ*L. The plate was incubated for 960 min at 37^°^C with 700 rpm shaking in double-orbital mode. OD was monitored over time every 13 min.

### Preparation of antibiotic-loaded GUVs

Giant unilamellar vesicles (GUVs) with encapsulated ciprofloxacin were prepared using the emulsion-transfer method [47, 54, 84].

A glass vial (1.5 mL) was loaded with 500 *µ*L of paraffin oil (Sigma-Aldrich) and 100 *µ*L of a 10 mg mL^−1^ solution of 1,2-dioleoyl-sn-glycero-3-phosphocholine lipids (DOPC, Avanti Polar Lipids) in chloroform. Afterwards, the solution was vortexed for 1 min, incubated for 60 min at ∼ 85^°^C to evaporate chloroform, and left for 15 min to cool down to room temperature. In the next step, 250 *µ*L of the lipid-oil solution (final lipid concentration 2 mg mL^−1^) were pipetted into a clean 1.5 mL glass vial and mixed with 25 *µ*L of the Inside-solution (I-solution, 3.2 *µ*g mL^−1^ ciprofloxacin and 100 *µ*M dextran (50 kDa, Sigma-Aldrich) in PBM9 (glucose replaced with 0.333% w/v sucrose)) by vortexing for ∼ 1 min. The generated turbid emulsion was layered on top of 150 *µ*L of the Outsisde-solution (O-solution, PBM9 (0.175% w/v glucose)) in a 1.5 mL DNase free Eppendorf tube, and centrifuged at 9000 *g* for 30 min. After centrifugation, first the oil and then the supernatant were removed from the sample, leaving 50 *µ*L of O-solution containing pelleted GUVs. In the final step, the resulting solution was diluted and resuspended by adding 100 *µ*L of fresh O-solution and gently pipetting it up and down 7-10 times.

### Assessment of antimicrobial responses in synthetic cell signalling network interfaced with bacteria

To test the response of the full netosis-like pathway (Figs 4**b, c** and Supplementary Fig. 16), four wells in a 96-well plate were loaded with 10 *µ*L of *E. coli* solution in PB(2×)M9 (*OD* = 0.66 before concentrating the cells 4×), prepared according to the aforementioned protocol, and 90 *µ*L of PB(2×)M9. All the wells were then topped with 100 *µ*L of PBS solution containing responsive or nonresponsive particles (*t*_*g*_ = 15 s, two wells with each particle type). Finally, 40 *µ*L of ciprofloxacin-loaded GUVs in PBM9 were added to two wells, each carrying different particle type, while the remaining two wells were filled with an identical volume of PBM9. The overall glucose concentration in all the samples was 0.175% w/v. Prepared samples were incubated in the plate reader described above for 1440 min at 37^°^C, with shaking at 500 rpm for 30 s every 5 min. OD was measured at 5 min intervals over time.

Prior to, and after incubation, all the samples were imaged with the aforementioned bright field and epifluorescence microscopy setup. Signals from core motifs (Alexa Fluor 594) and *E. coli* (EGFP) were recorded using green LED illumination/Texas Red filter and blue LED/GFP filter, respectively.

## ACKNOWLEDGMENTS

MW acknowledges support from the Engineering and Physical Sciences Research Council (EPSRC), the Department of Physics at the University of Cambridge (McLatchie Trust fund), and the Cambridge Philosophical Society. LDM acknowledges funding from a Royal Society University Research Fellowship (UF160152, URF \R \221009) and from the European Research Council (ERC) under the Horizon 2020 Research and Innovation Programme (ERC-STG No 851667 NANOCELL). LM, JK and PC acknowledge the UKRI grant EP/T002778/1. FR acknowledges funding from the Leverhulme Trust (EP/S023518/1). The authors thank Zoe Waller for useful discussions. A dataset underpinning these results is freely available at DOI: 10.17863/CAM.95810.

## AUTHOR CONTRIBUTIONS

MW designed the DNA sequences. MW and LM developed the experimental protocols. JK designed and manufactured the setup for efficient fabrication of coreshell particles. MW conducted all the experiments apart from CD of pH-responsive/nonresponsive DNA structures (FR), OD and pH measurements during bacterial growth in PBM9 medium (LM, JX), and ciprofloxacin MIC in *E. coli* measurement (LM), supported by LM. MW analyzed all the data. MW and LDM wrote the paper. MW, LM and LDM designed the research. LDM supervised the research, aided by PC. All authors discussed the results and edited the paper.

## Table of Contents

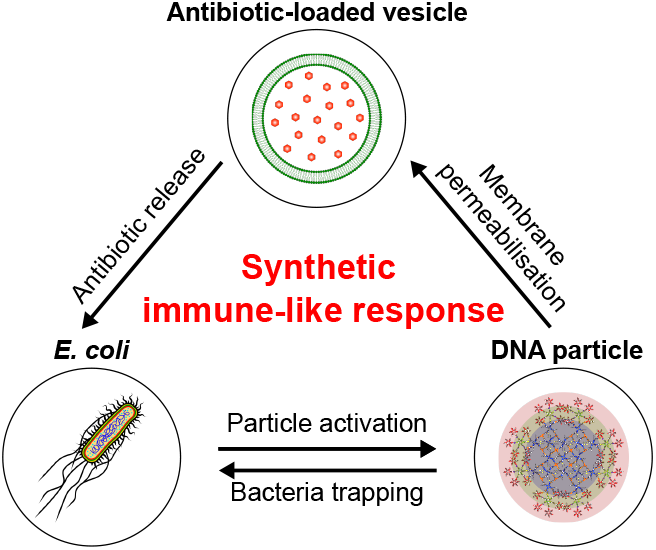

Bottom-up synthetic biology aims to replicate useful biological functions, but mimicking complex behaviours such as immunity remains elusive. Here, a synthetic consortium consisting of DNA particles and lipid vesicles is shown to detect bacteria, trap them, and expose them to antibiotics, imitating a core response of the innate immune system. These results demonstrate the bottom-up design of advanced life-like responses and outline new artificial-cell-based antimicrobial strategies.

## References

[1] Blain, J. C. & Szostak, J. W. Progress toward synthetic cells. Annu. Rev. Biochem. 83, 615–640 (2014).

[2] Buddingh’, B. C. & van Hest, J. C. M. Artificial Cells: Synthetic Compartments with Life-like Functionality and Adaptivity. Accounts of Chemical Research 50, 769–777 (2017).

[3] Cho, E. & Lu, Y. Compartmentalizing cell-free systems: Toward creating life-like artificial cells and beyond. ACS Synth. Biol. 9(11), 2881–2901 (2020).

[4] Zhu, Y., Guo, X., Liu, J., Li, F. & Yang, D. Emerging advances of cell-free systems toward artificial cells. Small Methods 4(10), 2000406 (2020).

[5] Boyd, M. A. & Kamat, N. P. Designing artificial cells towards a new generation of biosensors. Trends Biotechnol. 39(9), 927–939 (2021).

[6] Robinson, A. O., Venero, O. M. & Adamala, K. P. Toward synthetic life: Biomimetic synthetic cell communication. Curr. Opin. Chem. Biol. 64, 165–173 (2021).

[7] Slomovic, S., Pardee, K. & Collins, J. J. Synthetic biology devices for in vitro and in vivo diagnostics. Proc. Natl. Acad. Sci. U.S.A. 112(47), 14429–14435 (2015).

[8] Chen, Z. et al. Synthetic beta cells for fusion-mediated dynamic insulin secretion. Nat. Chem. Biol. 14, 86–93 (2018).

[9] Krinsky, N. et al. Synthetic cells synthesize therapeutic proteins inside tumors. Adv. Healthc. Mater. 7(1701163) (2018).

[10] Ding, Y., Contreras-Llano, L. E., Morris, E., Mao, M. & Tan, C. Minimizing context dependency of gene networks using artificial cells. ACS Appl. Mater. Interfaces 10(36), 30137–30146 (2018).

[11] Toparlak, O. D. et al. Artificial cells drive neural differentiation. Sci. Adv. 6(38) (2020).

[12] Salehi-Reyhani, A., Ces, O. & Elani, Y. Artificial cell mimics as simplified models for the study of cell biology. Exp. Biol. Med. 242(13), 1309–1317 (2017).

[13] Xu, C., Hu, S. & Chen, X. Artificial cells: from basic science to applications. Mater. Today 19(9), 516–532 (2016).

[14] Zhu, T. F. & Szostak, J. W. Coupled growth and division of model protocell membranes. J. Am. Chem. Soc. 131(15), 5705–5713 (2009).

[15] Kretschmer, S., Ganzinger, K. A., Franquelim, H. G. & Schwille, P. Synthetic cell division via membrane-transforming molecular assemblies. BMC Biol. 17(43) (2019).

[16] Steinkühler, J. et al. Controlled division of cell-sized vesicles by low densities of membrane-bound proteins. Nat. Commun. 11(905) (2020).

[17] Lee, K. Y. et al. Photosynthetic artificial organelles sustain and control ATP-dependent reactions in a protocellular system. Nat. Biotechnol. 36, 530–535 (2018).

[18] Berhanu, S., Ueda, T. & Kuruma, Y. Artificial photosynthetic cell producing energy for protein synthesis. Nat. Commun. 10(1325) (2019).

[19] Deshpande, S., Wunnava, S., Hueting, D. & Dekker, C. Membrane tension–mediated growth of liposomes. Small 15(38), 1902898 (2019).

[20] Downs, F. G. et al. Multi-responsive hydrogel structures from patterned droplet networks. Nat. Chem. 12, 363– 371 (2020).

[21] Zhang, S. et al. Engineering motile aqueous phase-separated droplets via liposome stabilisation. Nat. Commun. 12(1673) (2021).

[22] Mukwaya, V., Mann, S. & Dou, H. Chemical communication at the synthetic cell/living cell interface. Commun. Chem. 4(161) (2021).

[23] Smith, J. M., Chowdhry, R. & Booth, M. J. Controlling synthetic cell-cell communication. Front. Mol. Biosci. 8(809945) (2022).

[24] Garamella, J., Majumder, S., Liu, A. P. & Noireaux, V. An adaptive synthetic cell based on mechanosensing, biosensing, and inducible gene circuits. ACS Synth. Biol. 8(8), 1913–1920 (2019).

[25] Kaufhold, W. T., Brady, R. A., Tuffnell, J. M., Cicuta, P. & Di Michele, L. Membrane Scaffolds Enhance the Responsiveness and Stability of DNA-Based Sensing Circuits. Bioconjug. Chem. (2019).

[26] Samanta, A., Sabatino, V., Ward, T. R. & Walther, A. Functional and morphological adaptation in DNA protocells via signal processing prompted by artificial metalloenzymes. Nat. Nanotechnol. 15, 914–921 (2020).

[27] Leathers, A. et al. Reaction–diffusion patterning of DNA-based artificial cells. J. Am. Chem. Soc. 144(38), 17468– 17476 (2022).

[28] Rubio-Sánchez, R., Barker, S. E., Walczak, M., Cicuta, P. & Di Michele, L. A Modular, Dynamic, DNA-Based Platform for Regulating Cargo Distribution and Transport between Lipid Domains. Nano Lett. 21, 2800–2808 (2021).

[29] Elani, Y., Law, R. V. & Ces, O. Protein synthesis in artificial cells: using compartmentalisation for spatial organisation in vesicle bioreactor. Phys. Chem. Chem. Phys. 17, 15534–15537 (2015).

[30] van Nies, P. et al. Self-replication of DNA by its encoded proteins in liposome-based synthetic cells. Nat. Commun. 9(1583) (2018).

[31] Rideau, E., Dimova, R., Schwille, P., Wurm, F. R. & Landfeste, K. Liposomes and polymersomes: a comparative review towards cell mimicking. Chem. Soc. Rev. 47, 8572–8610 (2018).

[32] Stano, P. Gene expression inside liposomes: From early studies to current protocols. Chem. Eur. J. 25(33), 7798–7814 (2019).

[33] Niederholtmeyer, H., Chaggan, C. & Devaraj, N. K. Communication and quorum sensing in non-living mimics of eukaryotic cells. Nat. Commun. 9(5027) (2018).

[34] Huang, X. et al. Interfacial assembly of protein–polymer nano-conjugates into stimulus-responsive biomimetic protocells. Nat. Commun. 4(2239) (2013).

[35] Joesaar, A. et al. DNA-based communication in populations of synthetic protocells. Nat. Nanotechnol. 14, 369–378 (2019).

[36] Mason, A. F. & van Hest, J. C. M. Multifaceted cell mimicry in coacervate-based synthetic cells. Emerg. Top. Life Sci. 3(5), 567–571 (2019).

[37] Allen, M. E., Hindley, J. W., Baxani, D. K., Ces, O. & Elani, Y. Hydrogels as functional components in artificial cell systems. Nat. Rev. Chem. 6, 562–578 (2022).

[38] Xu, C., Martin, N., Li, M. & Mann, S. Living material assembly of bacteriogenic protocells. Nature 609, 1029– 1037 (2022).

[39] Rubio-Sánchez, R., Fabrini, G., Cicuta, P. & Di Michele, L. Amphiphilic DNA nanostructures for bottom-up synthetic biology. Chem. Commun. 57, 12725–127405 (2021).

[40] Marshall, J. S., Warrington, R., Watson, W. & Kim, H. L. An introduction to immunology and immunopathology. Allergy Asthma Clin. Immunol. 14(49) (2018).

[41] Diamond, M. S. & Kanneganti, T. Innate immunity: the first line of defense against SARS-CoV-2. Nat. Immunol. 23, 165–176 (2022).

[42] Lim, B. et al. Reprogramming synthetic cells for targeted cancer therapy. ACS Synth. Biol. 11(3), 1349–1360 (2022).

[43] Gardner, P. M., Winzer, K. & Davis, B. G. Sugar synthesis in a protocellular model leads to a cell signalling response in bacteria. Nat. Chem. 1, 377–383 (2009).

[44] Lentini, R. et al. Integrating artificial with natural cells to translate chemical messages that direct E. coli behaviour. Nat. Commun. 5(4012) (2014).

[45] Schwarz-Schilling, M., Aufinger, L., Mückl, A. & Simmel, F. C. Chemical communication between bacteria and cell-free gene expression systems within linear chains of emulsion droplets. Integr. Biol. 8(4), 564–570 (2016).

[46] Lentini, R. et al. Two-way chemical communication between artificial and natural cells. ACS Cent. Sci. 3(2), 117–123 (2017).

[47] Rampioni, G. et al. Synthetic cells produce a quorum sensing chemical signal perceived by Pseudomonas aeruginosa. Chem. Commun. 54, 2090–2093 (2018).

[48] Wang, X. et al. Chemical communication in spatially organized protocell colonies and protocell/living cell microarrays. Chem. Sci. 10(41), 9446–9453 (2019).

[49] Smith, J. M., Hartmann, D. & Booth, M. J. Engineering cellular communication between light-activated synthetic cells and bacteria. bioRxiv (2022).

[50] Gispert, I. et al. Stimuli-responsive vesicles as distributed artificial organelles for bacterial activation. Proceedings of the National Academy of Sciences 119(42), e2206563119 (2022).

[51] Brinkmann, V. et al. Neutrophil extracellular traps kill bacteria. Science 303(5663), 1532–1535 (2004).

[52] Kaplan, M. J. & Radic, M. Neutrophil extracellular traps: double-edged swords of innate immunity. J. Immunol. 189(6), 2689–2695 (2012).

[53] Papayannopoulos, V. Neutrophil extracellular traps in immunity and disease. Nat. Rev. Immunol. 18, 134–147 (2018).

[54] Walczak, M. et al. Responsive core-shell DNA particles trigger lipid-membrane disruption and bacteria entrapment. Nat. Commun. 12(4743) (2021).

[55] Sánchez-Clemente, R. et al. Study of pH changes in media during bacterial growth of several environmental strains. Proceedings 2(20), 1297 (2018).

[56] Shu, W. et al. DNA molecular motor driven micromechanical cantilever arrays. J. Am. Chem. Soc. 127(48), 17054–17060 (2005).

[57] Dvo?ráková, Z. et al. i-motif of cytosine-rich human telomere DNA fragments containing natural base lesions. Nucleic Acids Res. 46(4), 1624–1634 (2018).

[58] S?koláková, P. et al. Systematic investigation of sequence requirements for DNA i-motif formation. Nucleic Acids Res. 47(5), 2177–2189 (2019).

[59] Cerbino, R. & Trappe, V. Differential dynamic microscopy: Probing wave vector dependent dynamics with a microscope. Phys. Rev. Lett. 100(18), 188102 (2008).

[60] Cerbino, R. & Cicuta, P. Perspective: Differential dynamic microscopy extracts multi-scale activity in complex fluids and biological systems. J. Chem. Phys. 147, 110901 (2017).

[61] Brady, R. A., Brooks, N. J., Cicuta, P. & Di Michele, L. Crystallization of amphiphilic DNA c-stars. Nano Lett. 17(5), 3276–3281 (2017).

[62] Brady, R. A., Brooks, N. J., Fodera, V., Cicuta, P. & Di Michele, L. Amphiphilic-DNA platform for the design of crystalline frameworks with programmable structure and functionality. J. Am. Chem. Soc. 140(45), 15384– 15392 (2018).

[63] Brady, R. A., Kaufhold, W. T., Brooks, N. J., Fodera, V. & Di Michele, L. Flexibility defines structure in crystals of amphiphilic DNA nanostars. J. Phys.: Cond. Matter 31, 074003 (2019).

[64] Nesterova, I. V. & Nesterov, E. E. Rational design of highly responsive ph sensors based on dna i-motif. J. Am. Chem. Soc. 136(25), 8843–8846 (2014).

[65] Chen, N. et al. Real-time monitoring of dynamic microbial fe(iii) respiration metabolism with a living cell-compatible electron-sensing probe. Angew. Chem. 61(18), e202115572 (2022).

[66] Wang, L. et al. Engineering consortia by polymeric microbial swarmbots. Nature Communications 13, 3879 (2022).

[67] Balasubramanian, S., Aubin-Tam, M.-E. & Meyer, A. S. 3d printing for the fabrication of biofilm-based functional living materials. ACS Synthetic Biology 8, 1564–1567 (2019).

[68] Yamashita, T. & Yamamoto-Ikemoto, R. Nitrogen and phosphorus removal from wastewater treatment plant effluent via bacterial sulfate reduction in an anoxic bioreactor packed with wood and iron. Int. J. Environ. Res. Public Health 11(9), 9835–9853 (2014).

[69] Van Dillewijn, P., Nojiri, H., Van Der Meer, J. R. & Wood, T. K. Bioremediation, a broad perspective. Microb. Biotechnol. 2(2), 125–127 (2009).

[70] Rodrigo-Navarro, A., Sankaran, S., Dalby, M. J., del Campo, A. & Salmeron-Sanchez, M. Engineered living biomaterials. Nature Reviews Materials 6, 1175–1190 (2021).

[71] Ming, Z. et al. Living bacterial hydrogels for accelerated infected wound healing. Advanced Science 8, 2102545 (2021).

[72] Bhusari, S., Sankaran, S. & del Campo, A. Regulating bacterial behavior within hydrogels of tunable viscoelasticity. Advanced Science 9, 2106026 (2022).

[73] Hu, Y. et al. Identification of bacterial surface antigens by screening peptide phage libraries using whole bacteria cell-purified antisera. Front. Microbiol. 8(82) (2017).

[74] Memar, M. Y. & Baghi, H. B. Presepsin: A promising biomarker for the detection of bacterial infections. Biomed. Pharmacother. 111, 649–656 (2019).

[75] Mognetti, B. M., Cicuta, P. & Di Michele, L. Programmable interactions with biomimetic DNA linkers at fluid membranes and interfaces. Rep. Prog. Phys. 82, 116601 (2019).

[76] Morzy, D. et al. Cations regulate membrane-attachment and functionality of DNA nanostructures. J. Am. Chem. Soc. 143, 7358–7367 (2021).

[77] Zadeh, J. N. et al. NUPACK: analysis and design of nucleic acid systems. J. Comput. Chem. 32, 170–173 (2011).

[78] Mergny, J. L. & Lacroix, L. Analysis of thermal melting curves. Oligonucleotides 13(6), 515–537 (2003).

[79] Lord, N. Fluctuation timescales in bacterial gene expression. Ph.D. thesis, Harvard University (2014).

[80] Mancini, L. et al. A general workflow for characterization of nernstian dyes and their effects on bacterial physiology. Biophys. J. 118(1), 4–14 (2020).

[81] Bright, G. R., Fisher, G. W., Rogowska, J. & Taylor, D. L. Fluorescence ratio imaging microscopy: temporal and spatial measurements of cytoplasmic pH. J. Cell Biol. 104(4), 1019–1033 (1987).

[82] Surre, J. et al. Strong increase in the autofluorescence of cells signals struggle for survival. Sci. Rep. 8(12088) (2018).

[83] Wiegand, I., Hilpert, K. & Hancock, R. E. W. Agar and broth dilution methods to determine the minimal inhibitory concentration (MIC) of antimicrobial substances. Nat. Protoc. 3, 163–175 (2011).

[84] Fujii, S. et al. Liposome display for in vitro selection and evolution of membrane proteins. Nat. Protoc. 9, 1578– 1591 (2014).

