## Supplementary Information for "A synthetic signalling network imitating the action of immune cells in response to bacterial metabolism"

(Dated: April 15, 2023)

Supplementary Table 1. **Oligonucleotide sequences.** Labels: CM – core motif, ICM – inner corona motif, OCM – outer corona motif, CS – cholesterol strand, NCS – non-cholesterol strand, $\gamma$ - domain $\gamma$, $\gamma$* - domain $\gamma$*, $\delta$ - domain $\delta$, $\delta$* - domain $\delta$*, IM - domain $IM$, IM*-3m - domain $IM^* - 3m$, Rh6G - Rhodamine 6G, FLUO - Fluorescein, BHQ1 - Black Hole Quencher 1, A647 - Alexa Fluor 647, A594 - Alexa Fluor 594, TEG - Triethylene glycol.

| **Label** | **Oligonucleotide sequence (5' → 3')** |
|---:|---|
| CM.1 | GGAGGTGGAGGTAGTGGATGGCTCGTCACTGCATACCGGCTGCTCAGAACATACATAA CGCTTCGCCACGTCAAAC |
| CM.1 (IM) | GGAGGTGGAGGTAGTGGATGGCTCGTCACTGCATACCGGCTGCTCAGAACATACATAA CGCTTCCCAATCCCAATCCCAATCCC |
| CM.1 (IM, Rh6G) | GGAGGTGGAGGTAGTGGATGGCTCGTCACTGCATACCGGCTGCTCAGAACATACATAA CGCT Rhodamine 6G CCCAATCCCAATCCCAATCCC |
| CM.2 | GGAGGTGGAGGTAGTGGATGGCGTTATGTATGTTCTGAGCTCGGACTCGCAAACGCCA CGCTTTTTTTTTTTTTTTT |
| CM.2 (FLUO) | GGAGGTGGAGGTAGTGGATGGCGTTATGTATGTTCTGAGC Fluorescein CGGACTCGCA AACGCCACGC |
| CM.3 | GGAGGTGGAGGTAGTGGATGGCGTGGCGTTTGCGAGTCCGTCGTGTAAATGATGCGG TCGCTTCGCCACGTCAAAC |
| CM.3 (IM) | GGAGGTGGAGGTAGTGGATGGCGTGGCGTTTGCGAGTCCGTCGTGTAAATGATGCGG TCGCTTCCCAATCCCAATCCCAATCCC |
| CM.4 | GGAGGTGGAGGTAGTGGATGGCGACCGCATCATTTACACGTGCCGGTATGCAGTGACG AGCTTTTTTTTTTTTTTTT |
| CM.4 ($\gamma$) | GGAGGTGGAGGTAGTGGATGGCGACCGCATCATTTACACGTGCCGGTATGCAGTGACG AGCTTCGTCGGCGACGTCACG |
| ICM.1 | GCAATCGCCATCCGCGTCTGTTCGCATGATATCCGTGCTCCGGCTTCCAGCTCCG |
| ICM.2 | CGGAGCACGGATATCATGCGTTCGCCGTAGAGCAGCAATTCGGCTTCCAGCTCCG |
| ICM.3 | CGAATTGCTGCTCTACGGCGTTGGCCTCAGAGTCTGCGAACGGCTTCCAGCTCCG |
| ICM.4 | CGTTCGCAGACTCTGAGGCCTTCGGTATGAGCCTGTGCAACGGCTTCCAGCTCCG |
| ICM.5 | CGTTGCACAGGCTCATACCGTTCAATGGAGCCTTTACGGCACGCTTCCAGCTCCG |
| ICM.6 | GTGCCGTAAAGGCTCCATTGTTCAGACGCGGATGGCGATTGCGTTTGACGTGGCG |
| ICM.6 (IM*-3m) | GTGCCGTAAAGGCTCCATTGTTCAGACGCGGATGGCGATTGCTTGTGATTGGGATTTG GATTGTG |
| ICM.6 (IM*-3m, BHQ1) | GTGCCGTAAAGGCTCCATTGTTCAGACGCGGATGGCGATTGCTTGTGATTGGGATTTG GATTGTG Black Hole Quencher 1 |
| OCM.1 | GCAAGTGCTATGAGGTTGCGTTCCTGGTACTTTACATGTTCG |
| OCM.2 | CGAACATGTAAAGTACCAGGTTGCATCGATGGTCGACGAACG |
| OCM.3 | CGTTCGTCGACCATCGATGCTTGCCTTGACTGCCAATCAACG |
| OCM.3 (A647) | CGTTCGTCGACCATCGATGCTTGCCTTGACTGCCAATCAACG Alexa Fluor 647 |
| OCM.4 | CGTTGATTGGCAGTCAAGGCTTGCCGAAGTTGGAGCGGACCG |
| OCM.5 | CGGTCCGCTCCAACTTCGGCTTGTGAGCATCTATCTCTAAAC |
| OCM.6 | GTTTAGAGATAGATGCTCACTTCGCAACCTCATAGCACTTGCCGGAGCTGGAAGC |
| CS | CATCCACTACCTCCACCTCCAA TEG Cholesterol |
| NCS | CATCCACTACCTCCACCTCCAA |
| $\gamma$*+$\delta$* (A594) | Alexa Fluor 594 CGCACTTCCGGCTTGCCGTGACGTCGCCGACG |
| $\delta$ | GCAAGCCGGAAGTGCG |
| IM | CCCAATCCCAATCCCAATCCC |
| IM*-3m | GTGATTGGGATTTGGATTGTG |
| IM*-3m (BHQ1) | GTGATTGGGATTTGGATTGTG Black Hole Quencher 1 |

Supplementary Table 2. **Sample composition.** Labels as in Supplementary Table 1. For each strand used to prepare a specific sample, its concentration in $\mu$M is given in parentheses following the strand label. In parentheses where multiple concentrations are listed only one was used depending on the experiment (see figures and corresponding sections in Methods).

| **Sample** | **Strands** | **Relevant figures** |
|---|---|---|
| IM | IM (2/20) | 2d, 2e |
| IM*-3m | IM*-3m (2/20) | 2d, 2e |
| IM + IM*-3m | IM (2/20), IM*-3m (2/20) | 2d, 2e, S3 |
| CM | CM.1 (IM) (1), CM.2 (1), CM.3 (IM) (1), CM.4 (1), NCS (4) | S1, S2, S3 |
| ICM | ICM.1 (2), ICM.2 (2), ICM.3 (2), ICM.4 (2), ICM.5 (2), ICM.6 (IM*-3m) (2) | S1, S2 |
| OCM | OCM.1 (10), OCM.2 (10), OCM.3 (10), OCM.4 (10), OCM.5 (10), OCM.6 (10) | S1, S2 |
| CM + ICM | CM.1 (IM) (1), CM.2 (1), CM.3 (IM) (1), CM.4 (1), NCS (4), ICM.1 (2), ICM.2 (2), ICM.3 (2), ICM.4 (2), ICM.5 (2), ICM.6 (IM*-3m) (2) | S1, S2, S4 |
| ICM + OCM | ICM.1 (2), ICM.2 (2), ICM.3 (2), ICM.4 (2), ICM.5 (2), ICM.6 (IM*-3m) (2), OCM.1 (10), OCM.2 (10), OCM.3 (10), OCM.4 (10), OCM.5 (10), OCM.6 (10) | S1, S2 |
| CM + OCM | CM.1 (IM) (1), CM.2 (1), CM.3 (IM) (1), CM.4 (1), NCS (4), OCM.1 (10), OCM.2 (10), OCM.3 (10), OCM.4 (10), OCM.5 (10), OCM.6 (10) | S1, S2 |
| CM + ICM + OCM | CM.1 (IM) (1), CM.2 (1), CM.3 (IM) (1), CM.4 (1), NCS (4), ICM.1 (2), ICM.2 (2), ICM.3 (2), ICM.4 (2), ICM.5 (2), ICM.6 (IM*-3m) (2), OCM.1 (10), OCM.2 (10), OCM.3 (10), OCM.4 (10), OCM.5 (10), OCM.6 (10) | S1, S2 |
| Non-responsive particles (A594) | CM.1 (1), CM.2 (1), CM.3 (1), CM.4 ($\gamma$) (1), CS (4), ICM.1 (2), ICM.2 (2), ICM.3 (2), ICM.4 (2), ICM.5 (2), ICM.6 (2), OCM.1 (10), OCM.2 (10), OCM.3 (10), OCM.4 (10), OCM.5 (10), OCM.6 (10), $\gamma$*+$\delta$* (A594) (1), $\delta$ (1) | 2g, 4b, 4c, S5, S15 |
| Responsive particles (FLUO, A647) | CM.1 (IM) (1), CM.2 (FLUO) (1), CM.3 (IM) (1), CM.4 (1), CS (4), ICM.1 (2), ICM.2 (2), ICM.3 (2), ICM.4 (2), ICM.5 (2), ICM.6 (IM*-3m) (2), OCM.1 (10), OCM.2 (10), OCM.3 (A647) (10), OCM.4 (10), OCM.5 (10), OCM.6 (10) | 2b |
| Responsive particles | CM.1 (IM) (1), CM.2 (1), CM.3 (IM) (1), CM.4 (1), CS (4), ICM.1 (2), ICM.2 (2), ICM.3 (2), ICM.4 (2), ICM.5 (2), ICM.6 (IM*-3m) (2), OCM.1 (10), OCM.2 (10), OCM.3 (10), OCM.4 (10), OCM.5 (10), OCM.6 (10) | 3f, S9 |
| Responsive particles (A594) | CM.1 (IM) (1), CM.2 (1), CM.3 (IM) (1), CM.4 ($\gamma$) (1), CS (4), ICM.1 (2), ICM.2 (2), ICM.3 (2), ICM.4 (2), ICM.5 (2), ICM.6 (IM*-3m) (2), OCM.1 (10), OCM.2 (10), OCM.3 (10), OCM.4 (10), OCM.5 (10), OCM.6 (10), $\gamma$*+$\delta$* (A594) (1), $\delta$ (1) | 2g, 3d, 3e, 4b, 4c, S5, S15 |
| Responsive particles (FLUO) | CM.1 (IM) (1), CM.2 (FLUO) (1), CM.3 (IM) (1), CM.4 (1), CS (4), ICM.1 (2), ICM.2 (2), ICM.3 (2), ICM.4 (2), ICM.5 (2), ICM.6 (IM*-3m) (2), OCM.1 (10), OCM.2 (10), OCM.3 (10), OCM.4 (10), OCM.5 (10), OCM.6 (10) | S11, S12 |
| Responsive particles (Rh6G, BHQ1) | CM.1 (IM, Rh6G) (1), CM.2 (1), CM.3 (IM) (1), CM.4 (1), CS (4), ICM.1 (2), ICM.2 (2), ICM.3 (2), ICM.4 (2), ICM.5 (2), ICM.6 (IM*-3m, BHQ1) (2), OCM.1 (10), OCM.2 (10), OCM.3 (10), OCM.4 (10), OCM.5 (10), OCM.6 (10), added after preparation - IM*-3m (BHQ1) (2) | S13 |
| Responsive particles (Rh6G) | CM.1 (IM, Rh6G) (1), CM.2 (1), CM.3 (IM) (1), CM.4 (1), CS (4), ICM.1 (2), ICM.2 (2), ICM.3 (2), ICM.4 (2), ICM.5 (2), ICM.6 (IM*-3m) (2), OCM.1 (10), OCM.2 (10), OCM.3 (10), OCM.4 (10), OCM.5 (10), OCM.6 (10) | S13 |

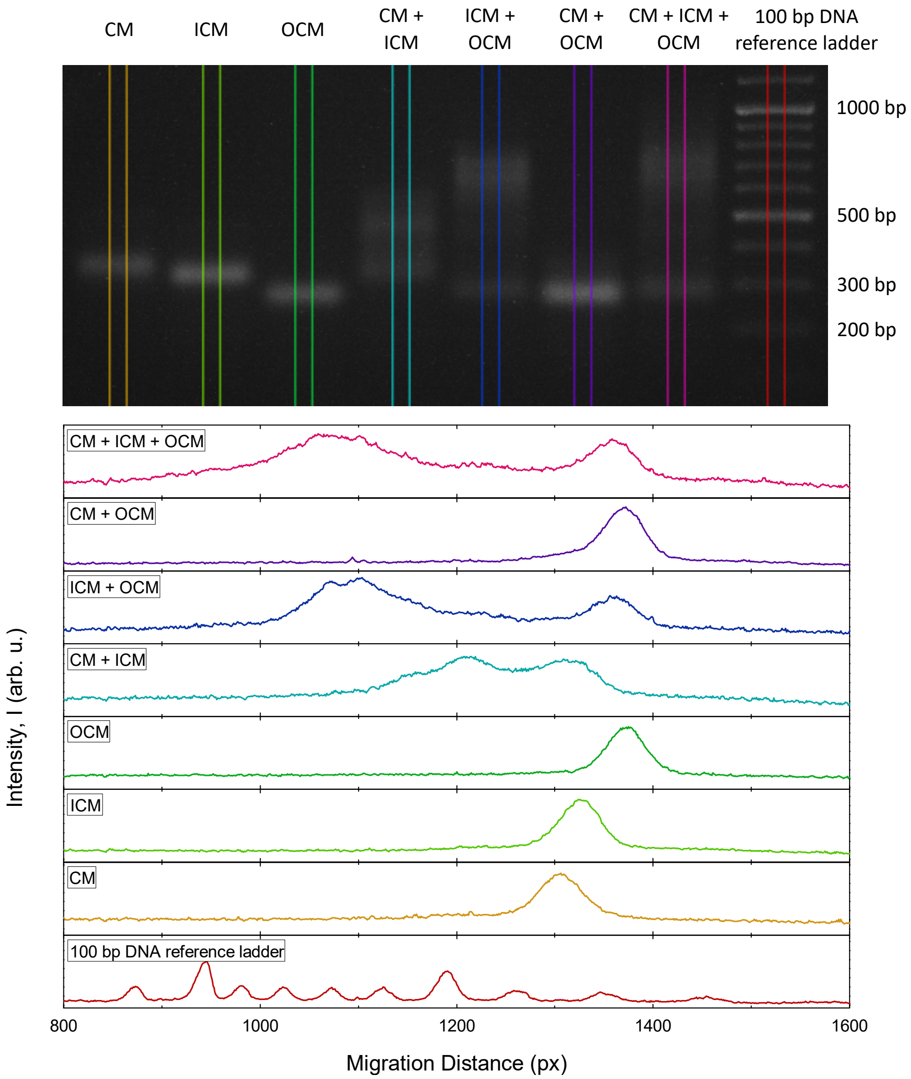


Supplementary Figure 1. **Agarose gel electrophoresis of non-functionalised DNA motifs.** All lanes with only one motif show an single bright band, assumed to contain DNA structures folded as intended. In the CM + OCM sample, the OCM band is very visible due to this construct being 10× more concentrated compared to CM, while CM produces a comparatively very faint band. The absence of bands corresponding to larger CM-OCM complexes demonstrates the lack of non-specific binding between structures programmed to solely interact with ICM. All the remaining samples display multiple distinct bands or a single smeared band indicating the successful formation of larger complexes through self-assembly of individual motifs.

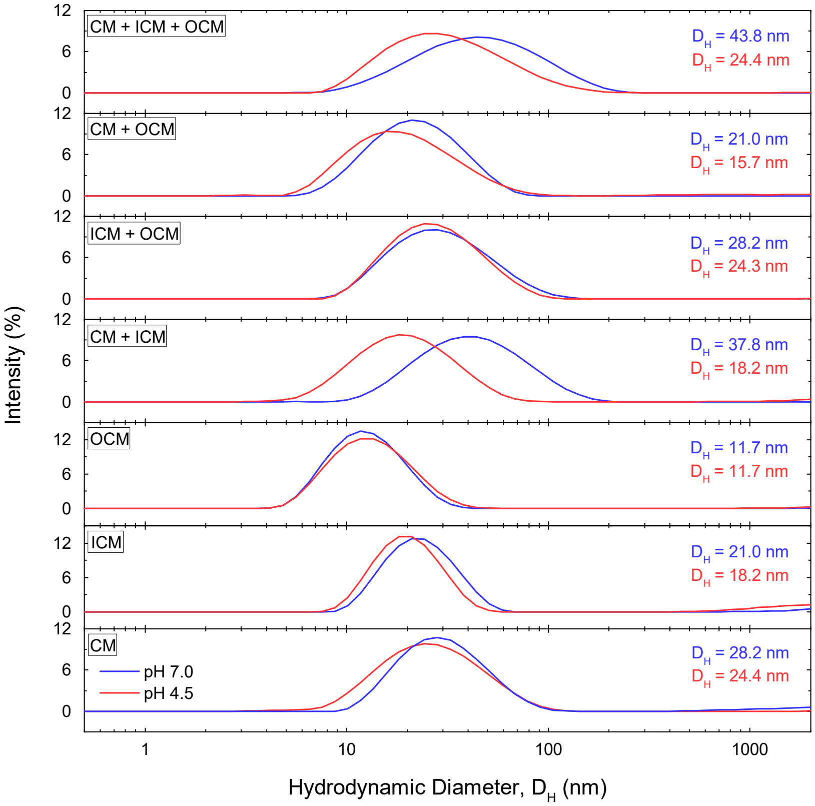


Supplementary Figure 2. **Dynamic light scattering of non-cholesterolised DNA nanostructures.** At neutral pH, an increase in $D_H$ is observed in samples featuring multiple motifs with complementary overhangs, indicating the successful formation of larger complexes (compare with Supplementary Fig. 1). Upon pH decrease below the transitional level required for i-motif formation (see Fig. 2**d**), a shift to smaller $D_H$ is detected in samples containing structures bridged by $IM\ /\ IM^{*}-3m$ bonds (CM + ICM and CM + ICM + OCM). This confirms i-motif formation and detachment of ICM from CM in response to buffer acidification. Data represent mean signal from six experiments conducted on two independently prepared samples (three per sample), each composed of fifteen measurements. $D_H$ was calculated by fitting each of the distinct bands to a lognormal distribution and extracting its maximum.

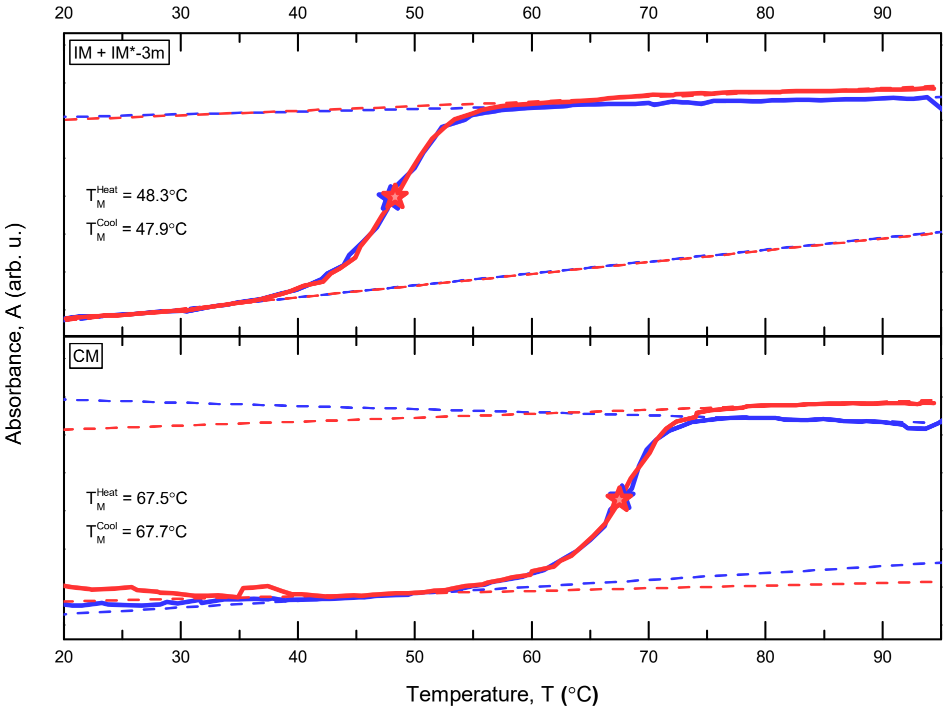


Supplementary Figure 3. **Experimental melting curves of core-forming DNA motifs and the corona-binding domain measured with UV-Vis spectroscopy.** The absorbance signal, recorded upon sample heating (red solid lines) and cooling (blue solid lines), displays a substantial separation ($\sim 20°$C) in the melting temperatures $T_M$ of core motifs (CM) and the domains $IM + IM^* - 3m$, bridging core motifs with inner corona motifs. This separation is required for the multi-step thermal annealing protocol reported in ref. [1] (also see Methods and main text). Data were collected in a single experiment. $T_M$ values (shown as stars) were extracted by fitting the high- and low-temperature plateaus of the absorbance curves with straight lines (dashed) and computing (*via* interpolation) the intersection between the data and the median of the two fits [2].

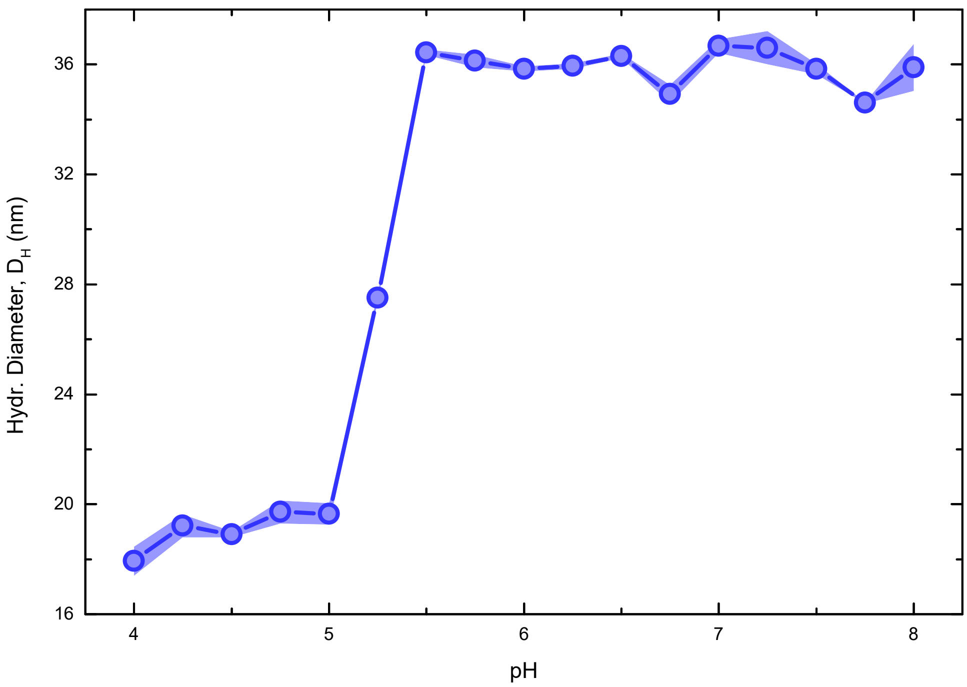


Supplementary Figure 4. **pH response curve of structures formed by cholesterol-free core and inner corona motifs obtained with DLS.** $D_H$ of core (CM) and inner corona (ICM) motif complexes shifts to smaller sizes once the pH is reduced below $\sim 5.50$ indicating i-motif formation and subsequent partial (pH $\sim 5.25$) or complete (pH $\sim 5.00$) detachment of ICM from CM. The transitional pH range observed here is comparable with its equivalent recorded by measuring UV absorbance at 297 nm [3, 4] (see Fig. 2**d**). Each experimental point represents the average value of $D_H \pm$ standard error (shaded regions) from six experiments of fifteen measurements each, conducted on two independently prepared samples. When not visible, errors are smaller than lines/symbols.

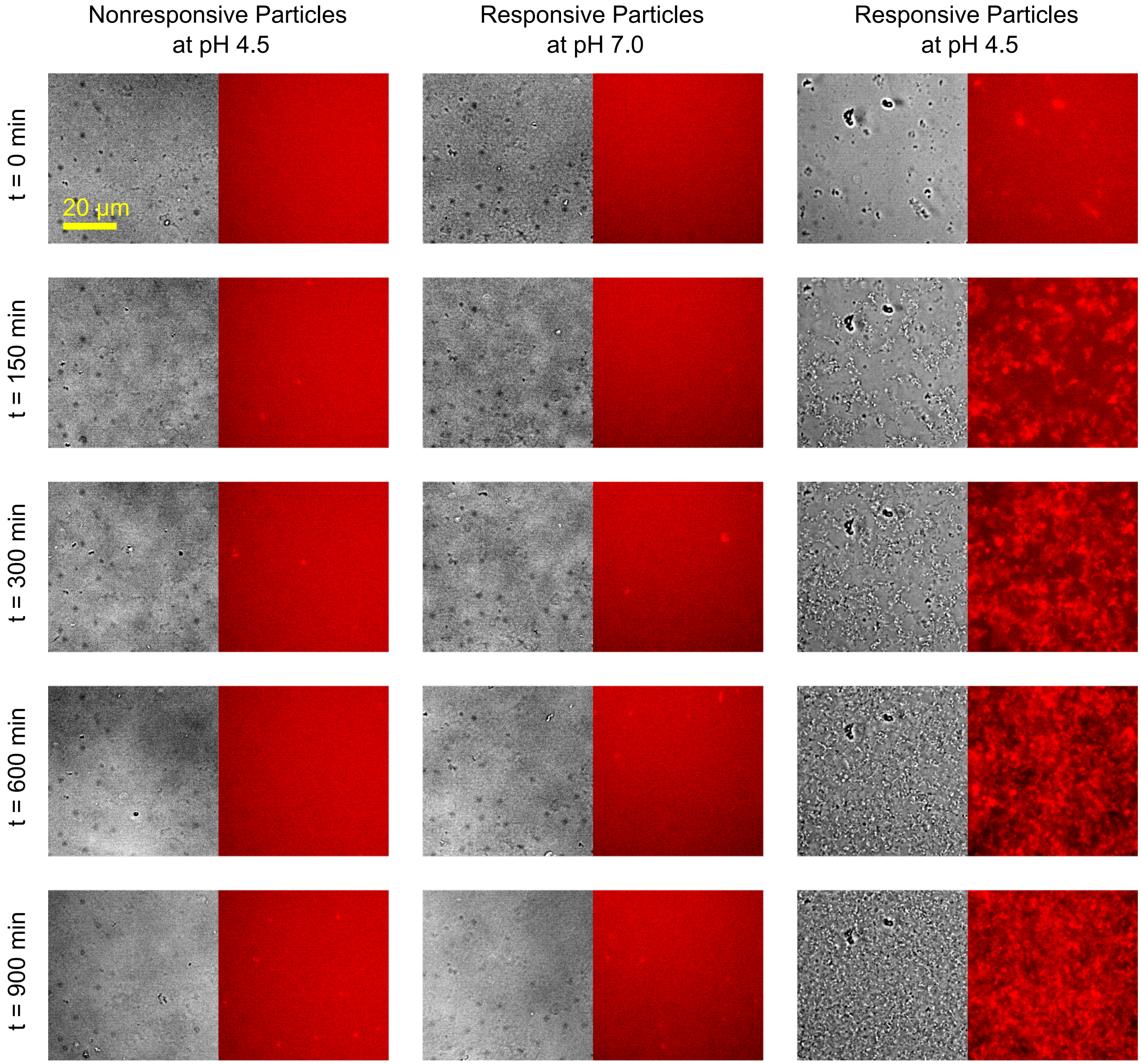


Supplementary Figure 5. **Bright field and epifluorescence micrographs illustrating particle aggregation triggered by pH decrease.** Progressive aggregation of cholesterol-rich particle cores triggered by buffer acidification can be observed in samples containing responsive particles in PBM9 medium. Control samples of responsive particles at neutral pH or nonresponsive particles at acidic pH, both in PBM9 medium, show no sign of aggregation. The hydrochloric acid (HCl), causing pH decrease, was added at time 0. Core motifs were labeled with Alexa Fluor 594 (red).

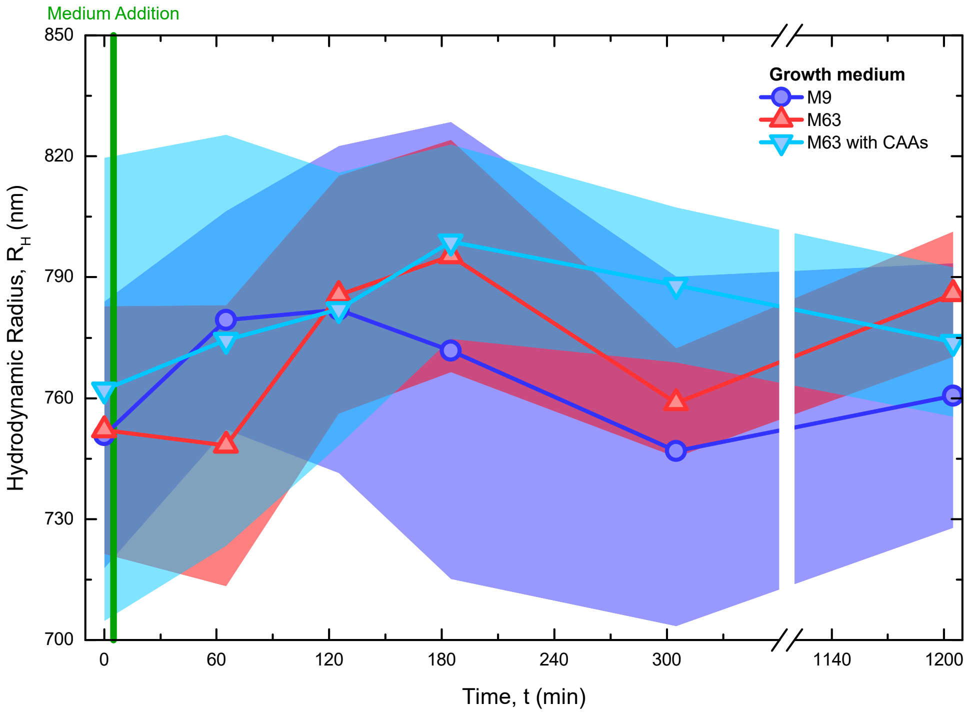


Supplementary Figure 6. **Particle stability in bacterial growth media as determined with differential dynamic microscopy (DDM).** After deployment in various bacterial growth media, including M9, M63 and M63 supplemented with casamino acids (CAAs, see Methods for the chemical composition of each medium), the particles ($t_g = 600$ s) remain stable as demonstarted by the lack of significant changes in their hydrodynamic radius obtained with DDM. For each sample, $R_H$ was measured for 1200 min following the growth medium addition at $t = 5$ min, with an additional measurement taken 5 min before adding the growth medium ($t = 0$). Data are shown as mean ± standard error (shaded regions) of three measurements conducted on three independently prepared samples.

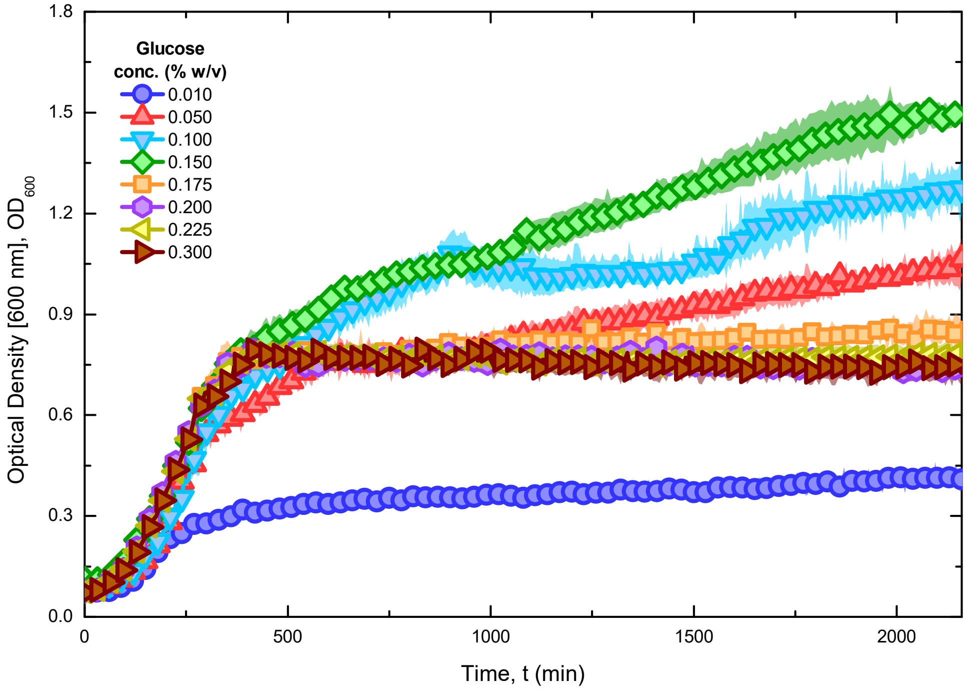


Supplementary Figure 7. **Bacterial growth in PBM9 medium with varying glucose concentration as determined *via* optical density measurements.** Greater *E. coli* yields, determined from optical density ($OD$), were achieved with intermediate glucose concentrations. Glucose concentrations above $\geq 0.175\%$ w/v were less favourable, possibly due to excessive medium acidification (see Fig. 3**b**, **f** and Supplementary Fig. 8). Data are shown as average $\pm$ standard error (shaded regions) of three measurements, each conducted on an independently prepared sample. When not visible, errors are smaller than the symbols.

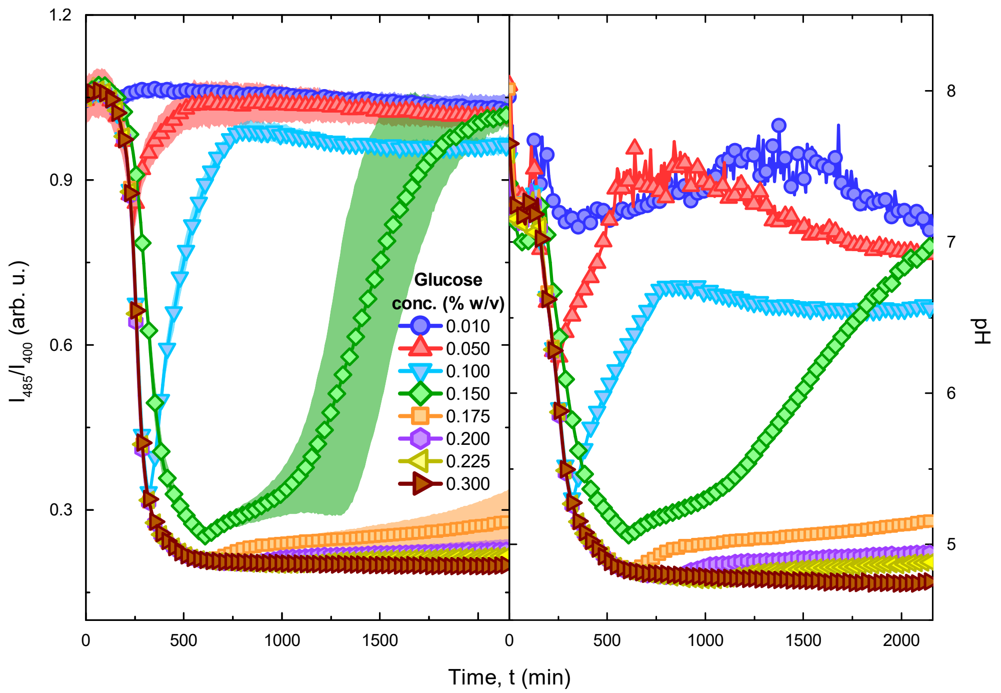


Supplementary Figure 8. **pH change accompanying bacterial growth in PBM9 medium with different glucose concentrations, tracked with the fluorimetric pH probe FITC-dextran.** Glucose metabolism in *E. coli* leads to pH decrease in solution (see Fig. 3**a**, **b**). Acquired data indicates that the magnitude of the pH change increases with glucose concentration. The pH increase observed in some samples at late times suggests re-uptake of acetate by the cells. The pH was calculated from the normalised fluorescence intensity ratio of FITC-dextran at excitation wavelengths of 485 nm and 400 nm ($I_{485}/I_{400}$), using the calibration curve shown in Supplementary Fig. 9. Each experimental point represents the average value of $I_{485}/I_{400}$ ± standard error (shaded regions) acquired from three independently prepared samples. When not visible, errors are smaller than lines/symbols.

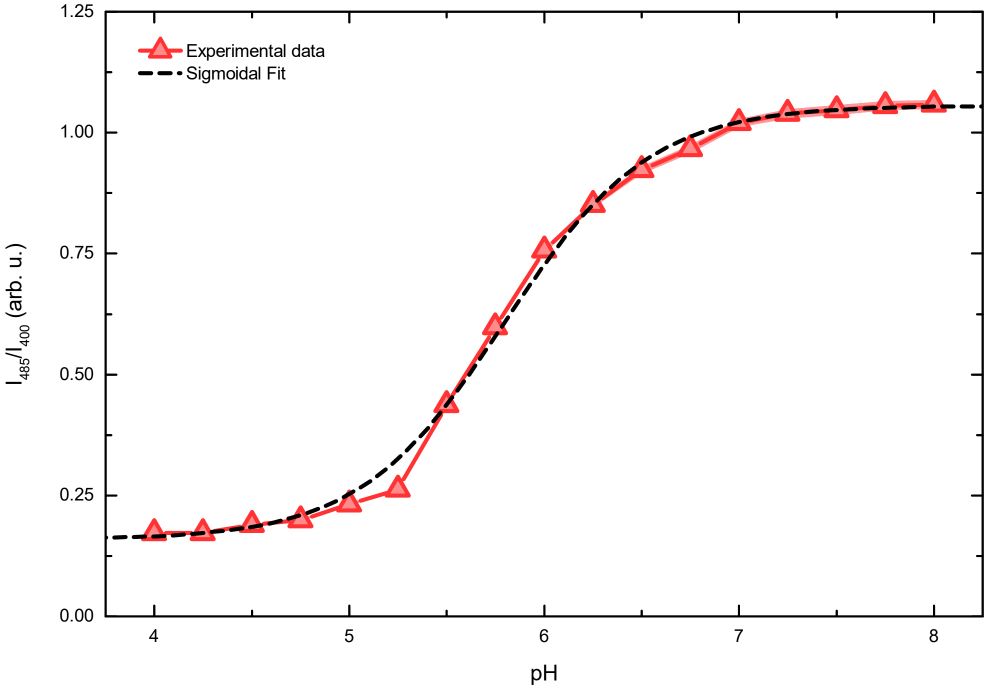


Supplementary Figure 9. **Calibration curve of the normalised fluorescence intensity ratio $I_{485}/I_{400}$ of a ratiometric pH probe FITC–dextran as a function of pH in PBM9 medium.** Experimental points were fitted to a sigmoid, and the resulting fit was used to calculate the pH from in Fig. **3f** and Supplementary Figs 8, 12. The obtained trend is in good agreement with the calibration performed on a similar fluorescein-based probe, reported in ref. [5]. Data represent the average ± standard error (shaded regions) of three measurements, each conducted on an independently prepared sample. When not visible, errors are smaller than lines/symbols.

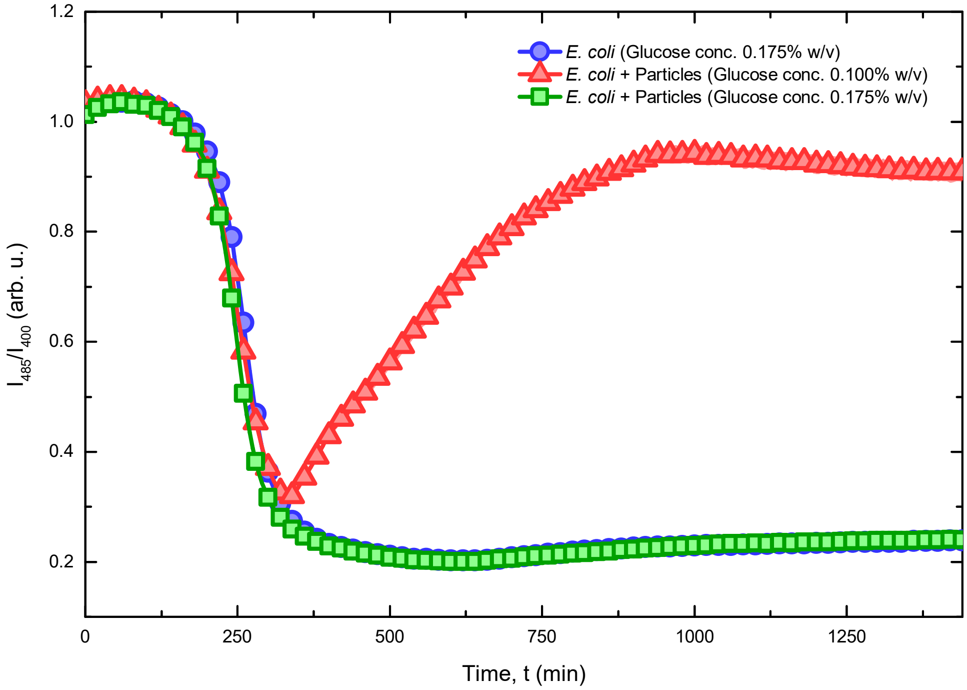


Supplementary Figure 10. **Normalised fluorescence intensity ratio $I_{485}/I_{400}$ of FITC-dextran recorded during an *E. coli* trapping experiment.** These data were used to produce the pH traces displayed in Fig. **3f**, using the calibration curve in Supplementary Fig. 9. Data are shown as the average ± standard error (shaded regions) of three measurements conducted on three independently prepared samples. When not visible, errors are smaller than lines/symbols.

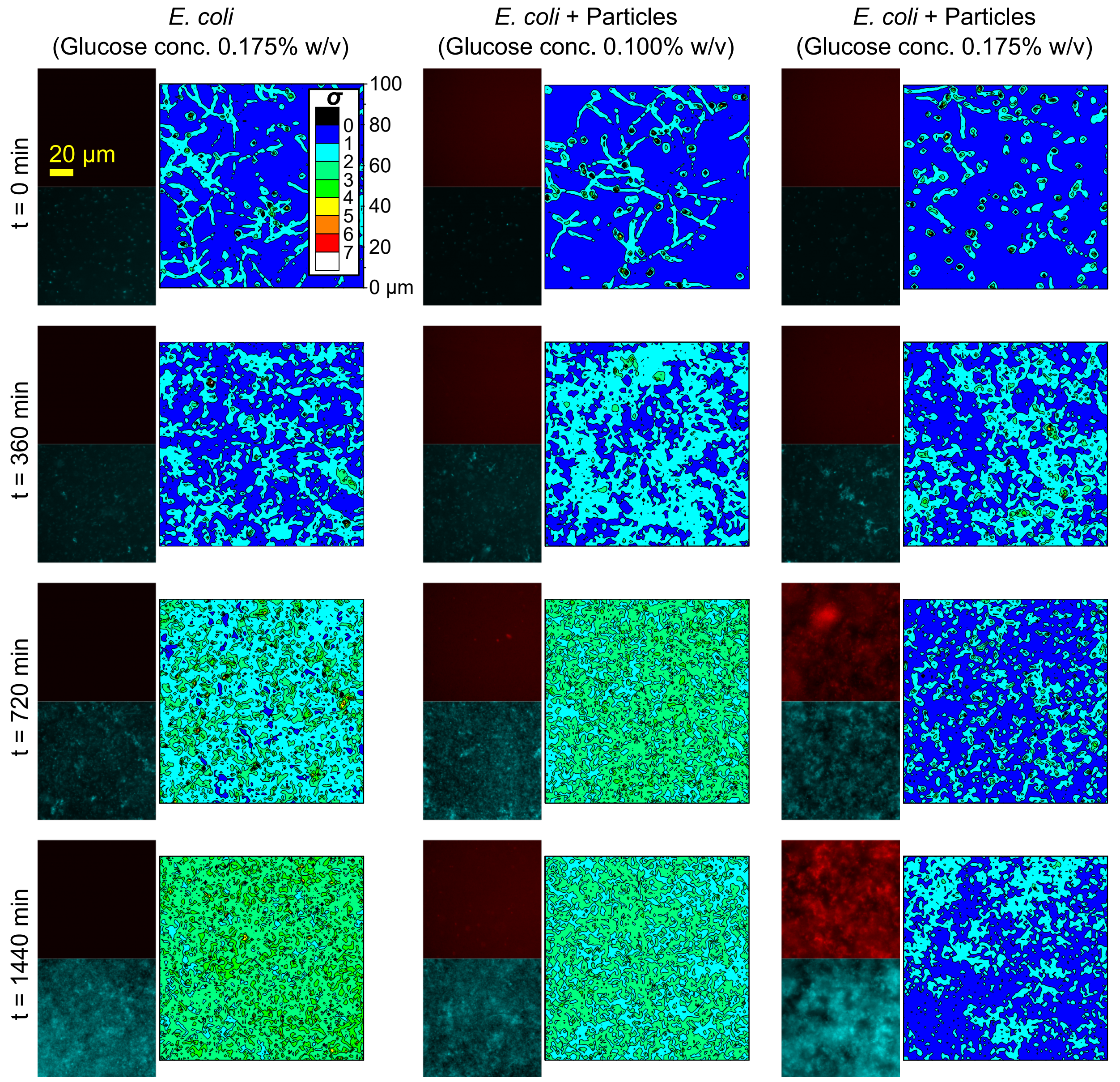


Supplementary Figure 11. **Epifluorescence micrographs and corresponding motility colormaps showing the progressive immobilisation of *E. coli* in a DNA network formed by particles activated by bacteria-induced medium acidification.** The colormaps illustrate the spatial distribution of the motility parameter $\sigma$ (see Methods) calculated from bright field videos for samples containing *E. coli* and responsive particles, with glucose concentration inhibiting (0.100% w/v) or supporting (0.175% w/v) particle activation, and a control sample with *E. coli* but lacking particles (glucose concentration = 0.175% w/v). An initial increase of $\sigma$ was observed in all the samples, resulting from bacterial growth (see Fig. 3**b**, **f** and Supplementary Fig. 7). Once the pH was sufficiently low to activate particle aggregation, *E. coli* became trapped in the assembled DNA network, which can be observed through stabilisation of $\sigma$ values. In the absence of particles, or in conditions preventing their activation, the parameter $\sigma$ continued to grow progressively. In epifluorescence micrographs, core motifs are shown in red (Alexa Fluor 594), *E. coli* in cyan (EGFP).

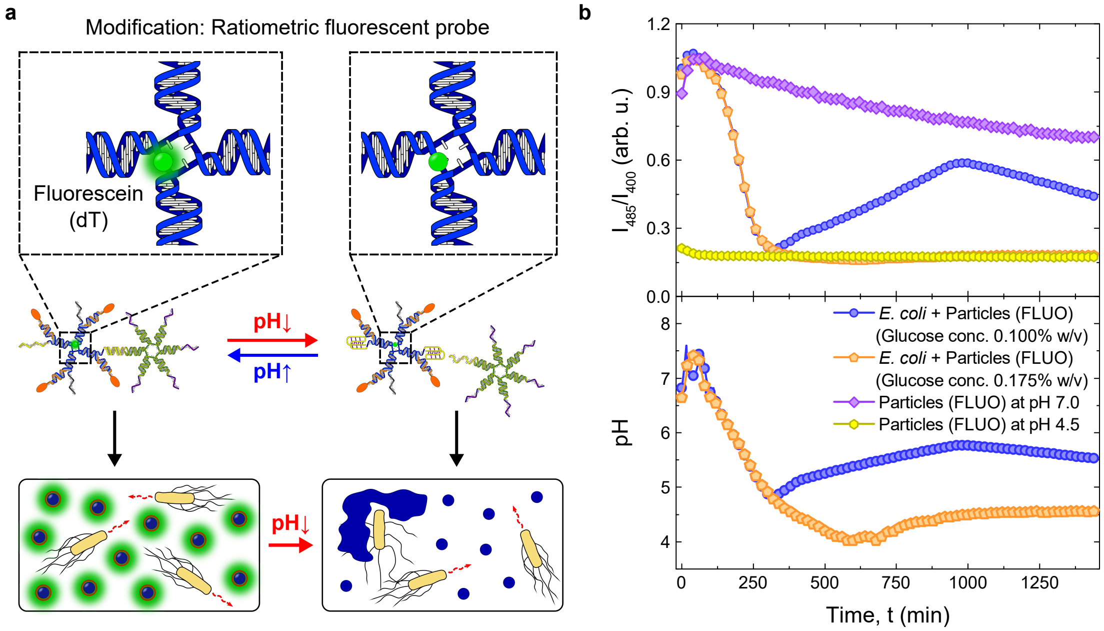


Supplementary Figure 12. ***E. coli* sensing with DNA particles functionalised with a fluorescent pH probe.** **a.** Diagram illustrating the mechanism of bacterial detection based on fluorescein-functionalised DNA particles. Acidification of the growth medium caused by glucose metabolism in *E. coli* affects emission of fluorescein fluorophores (green) located at the internal junction of core motifs (blue), leading to the progressive decrease the ratio between intensities recorded upon excitation at 485 nm and 400 nm. **b.** Time-trace of the fluorescence intensity ratio $I_{485}/I_{400}$ of fluorescein (FLUO) in PBM9 medium containing *E. coli*, fluorescein-functionalised particles and different glucose concentrations (top). The intensity ratio can be used to determine pH, calculated from a calibration curve shown in Supplementary Fig. 9, and track medium acidification induced by glucose metabolism (bottom). For bacteria containing samples, $I_{485}/I_{400}$ values are shown as the average ± standard error (shaded regions) of three measurements, each conducted on an independently prepared sample. When not visible, errors are smaller than lines/symbols.

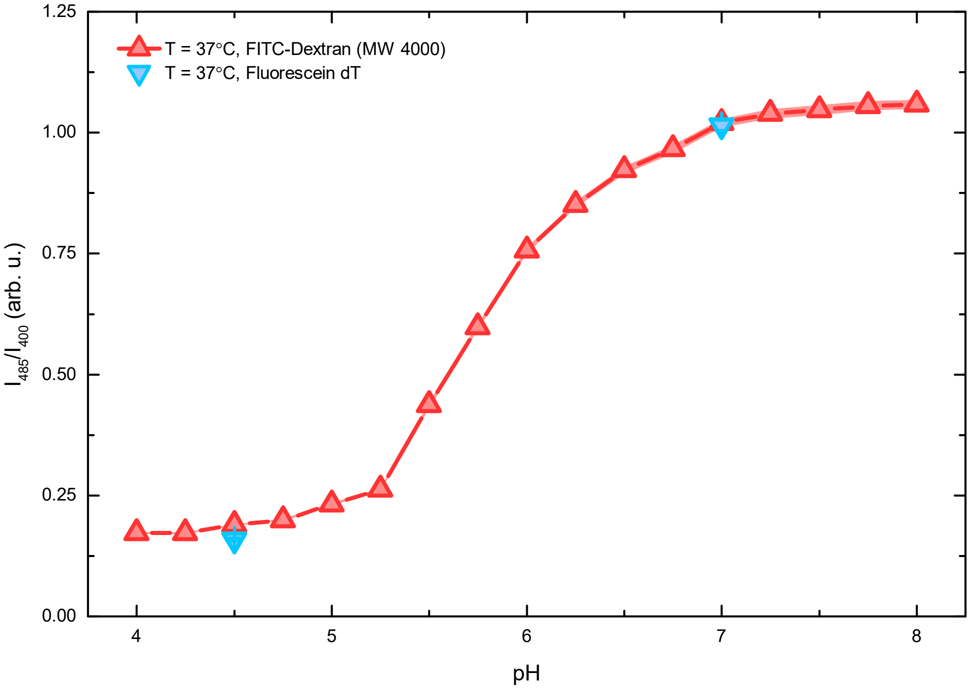


Supplementary Figure 13. **Comparison of the normalised fluorescence intensity ratios $I_{485}/I_{400}$ measured for fluorescein-based pH probes linked to dextran or to the DNA particles (dT).** Recorded $I_{485}/I_{400}$ values are nearly identical in the two systems, indicating that the calibration curve obtained for FITC-dextran (see Supplementary Fig. 9) can be used with fluorescein-functionalised DNA particles. Data are shown as mean ± standard error (shaded regions) of three measurements conducted on three independently prepared samples. When not visible, errors are smaller than lines/symbols.

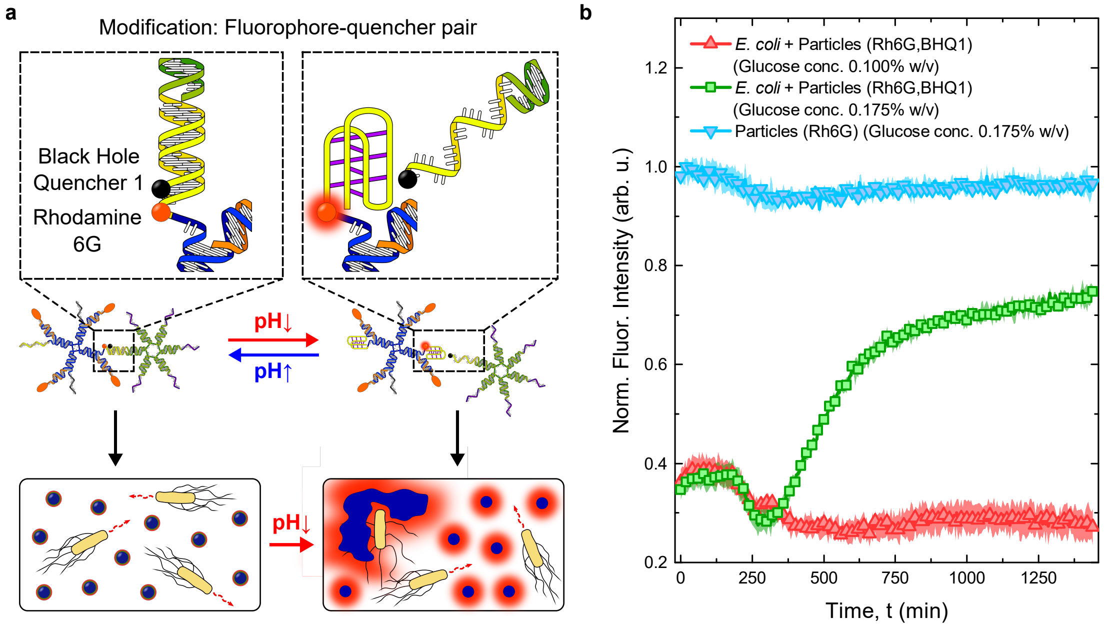


Supplementary Figure 14. **Detection of bacteria-induced acidification using fluorophore-quencher modified DNA particles.** **a.** Diagram showing the mechanism of bacterial detection based on pH-responsive particles functionalised with a fluorophore-quencher pair. Bacteria-induced corona detachment from particles leads to separation of a fluorophore (Rhodamine 6G, orange) and a quencher (Black Hole Quencher 1, black), resulting in increased emission by Rhodamine 6G. **b.** Time-trace of the normalised fluorescence intensity of Rhodamine 6G (Rh6G) for samples containing *E. coli* and fluorophore-quencher labelled particles with glucose concentration preventing (0.100% w/v) or triggering (0.175% w/v) particle activation, and a control sample with responsive particles lacking the quencher. The emission increase seen in fluorophore-quencher labelled particles and high-glucose concentration confirms the ability of the system to detect bacteria-induced medium acidification. Data are shown as the average ± standard error (shaded regions). of three measurements, each conducted on an independently prepared sample. When not visible, errors are smaller than lines/symbols.

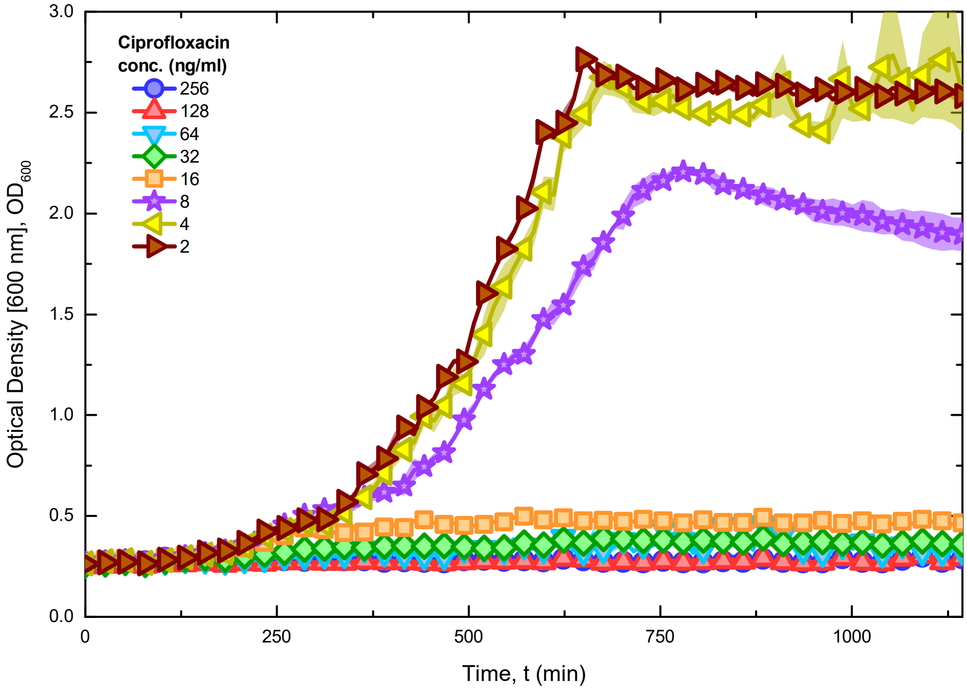


Supplementary Figure 15. **Minimum inhibitory concentration of the antibiotic ciprofloxacin for *E. coli* as determined *via* turbidity measurements.** The growth of *E. coli* in PBM9 medium was suppressed in the presence of ciprofloxacin at concentrations $\geq 16$ ng mL$^{-1}$. Consequently, the minimum inhibitory concentration (MIC) of ciprofloxacin for *E. coli* is estimated to be 16 ng mL$^{-1}$. Data represent the average value of $OD$ $\pm$ standard error (shaded regions) obtained from three independently prepared samples. When not visible, errors are smaller than lines/symbols.

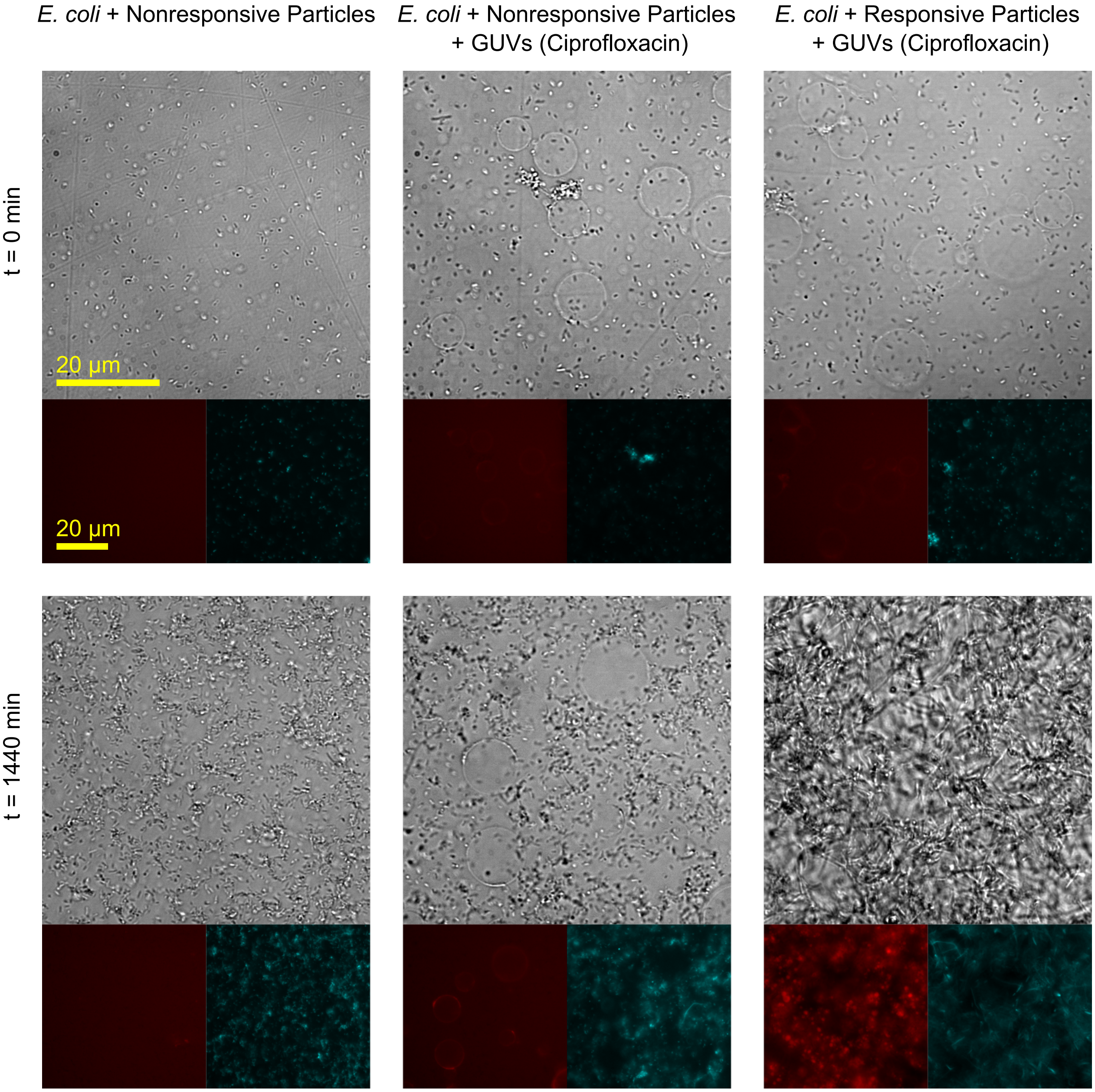


Supplementary Figure 16. **Bright field and epifluorescence micrographs demonstrating bacteria elongation caused by antibiotic release from vesicles destabilised by triggered DNA particles.** A substantial increase in *E. coli* length and disappearance of giant unilamellar vesicles (GUVs) was observed in samples containing all the components required to support the netosis-like response. These results indicate a successful rupture of GUVs induced by DNA aggregation, followed by ciprofloxacin release responsible for *E. coli* elongation, a consequence of their inability to divide. In the absence of GUVs or after replacing particles with their nonresponsive counterparts, no deformation of bacteria cells was detected. Core motifs are labelled with Alexa Fluor 594 (red), *E. coli* express EGFP (cyan).

*

[1] Walczak, M. *et al.* Responsive core-shell DNA particles trigger lipid-membrane disruption and bacteria entrapment. *Nat. Commun.* **12(4743)** (2021).

[2] Mergny, J. L. & Lacroix, L. Analysis of thermal melting curves. *Oligonucleotides* **13(6)**, 515–537 (2003).

[3] Dvořáková, Z. *et al.* i-Motif of cytosine-rich human telomere DNA fragments containing natural base lesions. *Nucleic Acids Res.* **46(4)**, 1624–1634 (2018).

[4] Školáková, P. *et al.* Systematic investigation of sequence requirements for DNA i-motif formation. *Nucleic Acids Res.* **47(5)**, 2177–2189 (2019).

[5] Bright, G. R., Fisher, G. W., Rogowska, J. & Taylor, D. L. Fluorescence ratio imaging microscopy: temporal and spatial measurements of cytoplasmic pH. *J. Cell Biol.* **104(4)**, 1019–1033 (1987).
